# FvM4K1, a Serine/Threonine Kinase and Ste20 Homolog Positively Regulates Fruit Size via Hippo Signalling Pathway in Woodland Strawberry (*Fragaria vesca*)

**DOI:** 10.1101/2024.12.17.628930

**Authors:** Si Gu, Xinghua Nie, Ling Qin, Yu Xing, Baoxiu Qi

## Abstract

Strawberry fruit size is critical for its marketability. However, organ size control is a complex process regulated by various signaling pathways. In animals, the Hippo signaling pathway acts as a negative regulator of organ size, with Ste20 kinase, a serine/threonine kinase being a key component. Mutation in Ste20 causes excessive cell proliferation and enlarged organs. In this study, FvM4K1, a Ste20-like kinase from woodland strawberry (*Fragaria vesca*), is identified. FvM4K1 partially restores defects in a yeast Ste20 mutant *ste20Δ* and fully rescues growth in Arabidopsis mutant *atsik1-4* lacking its Ste20 homolog, AtSIK1, underscoring its functional conservation. Downregulation FvM4K1 by RNAi in woodland strawberry leads to fruits and smaller plants resulting from reduced cell size and number, while overexpression increases organ size, indicating a positive role in fruit and organ size control which contrasts the negative role of Ste20 in other organisms. FvM4K1 auto-phosphorylates, with Lys269 and Thr396 being critical for its function. FvM4K1 interacts with FvMOB1A and FvMOB1B, components of the Hippo signalling pathway, and phosphorylates them at Thr35 and Thr36, respectively. These findings provide novel insights into the mechanisms underlying fruit and organ size control in woodland strawberry and contribute to our understanding of the Hippo signalling pathway in higher plants, a pathway that remains largely unexplored. It also opens new avenues for exploring the regulatory function of Hippo pathway in plant development and potentially inform biotechnological strategies for crop improvement.

## Introduction

Strawberry is a valuable crop and its fruit size is a key factor in determining its market value. Organ size control is a complex and highly coordinated process influenced by various signalling pathways. The Hippo pathway, initially discovered in fruit fly (*Drosophila melanogaster*), is a critical regulator of organ size in animals (Wu et al., 2003). Mutations in the Hippo signalling pathway can lead to uncontrolled cell proliferation and cancer (Han, 2019). However, its function in plants remains largely unknown.

Study in yeast (*Saccharomyces cerevisiae*) have shown that the serine/threonine kinase family Ste20 (Sterile 20) is a central component of the Hippo signalling pathway. Ste20 includes two subfamilies, the p21 activated kinases (PAKs) and the germinal centre kinases (GCKs), which differ in the location of their kinase domains: PAKs have a C-terminal, while GCKs have a N-terminal kinase domain (Boyce and Andrianopoulos, 2011). The yeast Ste20p is a typical member of the PAK subfamily and plays important roles including mating regulation (Wu et al., 1995), bud site selection (Sheu et al., 2000) and mitosis exit (Höfken and Schiebel, 2002) whilst Cdc15p of the GCKs is one of the core components in the mitotic exit network (Rock et al., 2013). Hippo (Hpo), a FCK and homolog of Cdc15p in flies, is involved in controlling organ size via proliferation and apoptosis (Wu et al., 2003). Flies have five core components in Hippo signalling pathway: Hpo, its scaffolding protein Salvador (Sav), the Nuclear Dbf2-related (NDR) kinase Warts (Wts), its helper Mps One Binder (MOB1) protein Mats, and the transcription factor Yorkie (Yki). Hpo, in association with Sav phosphorylates the downstream Wts/Mats complex. This is followed by phosphorylation and inactivation of the transcription factor Yki in the cytoplasm by 14-3-3 mediated ubiquitin degradation, resulting in growth inhibition (Zhao et al., 2011; Zheng and Pan, 2019). When Hippo signalling is inactivated, Yki remains unphosphorylated and translocates to the nucleus where it binds with Scalloped (sd) to activate the expression of target genes, including *cyclin-E* (*cycE*), *diap1* and *bantam*. This activation promotes proliferation and inhibits apoptosis, resulting in larger wings and other organs due to excessive cell growth and proliferation (Zhao et al., 2011; Zheng and Pan, 2019). In humans, the Ste20 kinase Mst1/2 interacts with and phosphorylates the scaffold protein WW45, leading to phosphorylation and activation of the downstream complex consisting of Lats1/2 and MOB1s. The transcription factor Yes-associated protein (YAP) and the transcriptional co-activator TAZ (PDZ-binding motif) are inactivated by phosphorylation by the LATS1/2 and MOB1 complex, promoting cell proliferation and cancer (Zheng and Pan, 2019). Recent advances in the understanding of the Hippo signalling pathway have led to the identification of more than 30 components (Zhao et al., 2011; Zheng and Pan, 2019).

Studies on the Hippo signalling pathway in plants are limited and primarily focused on the model plant Arabidopsis. Only a few components have been identified so far, including the Ste20 kinase AtSIK1 (Xiong et al., 2016), the MOB1 homologs AtMOB1A and AtMOB1B (Guo et al., 2020), and eight NDR homologs (Zhou et al., 2021). Mutants lacking *AtSIK1* exhibited inhibited growth characterized by reduced plant height, smaller leaves, and flowers, due to reduced cell size and number. This suggested that AtSIK1 plays a role in controlling organ size by regulating cell expansion and proliferation (Xiong et al., 2016). AtSIK1 interacts with AtMOB1A&B, homologues of MOB1 (Xiong et al., 2016). The double mutant *mob1a−/− 1b+/−* displayed a similar phenotype to *sik1−/−*, while the double mutant *sik1−/−mob1a+/−* was much smaller than the respective single mutants. In addition, the homozygous double mutant *sik1−/−mob1a−/−* struggled to survive (Guo et al., 2020). These findings suggest that AtSIK1 and AtMOB1s function in the same pathway and positively regulate organ size via the Hippo signalling pathway, different to the negative role of the Hippo signalling pathway in animals.

AtNDRs are primarily required for pollen development, fertilization, and seed set in Arabidopsis (Yoon et al., 2021). There are eight AtNDRs which appear to function redundantly, as single and double mutants of *atndr2*, *atndr4*, and *atndr5* show no noticeable phenotype. However, the *atndr2/atndr4/atndr5* triple mutant produces much shorter siliques with reduced fertility than wild type (Zhou et al., 2021). NDR2/4/5 interact with MOB1 proteins and regulate pollen development (Zhou et al., 2021). However, the interaction between AtSIK1 and AtNDRs has not yet been explored although interaction between the homologous proteins Ste20 and NDRs in yeast and animals are verified (Rock et al., 2013; Zhao et al., 2011; Zheng and Pan, 2019).

Therefore, the current understanding of the Hippo singling pathway in Arabidopsis suggests that it may function differently from its role in yeast, flies, and animals. While Hippo signaling in other organisms typically acts to inhibit cell proliferation and limit organ size, in Arabidopsis it appears to positively regulate cell proliferation and growth. Additionally, far fewer components in this pathway have been identified in Arabidopsis than in other species, and importantly, it remains unclear how these limited components coordinate cell proliferation and organ size. Validating the presence and function of homolog proteins in Hippo signalling pathway in another plant, especially in a crop species, could provide valuable insights into their role in organ size regulation, a key trait in plant breeding.

The aim of this study is to investigate the role of the Ste20 kinase homolog FvM4K1 in woodland strawberry, a model plant for the Rosaceae family. Through complementation studies in both yeast and Arabidopsis FvM4K1 was confirmed as a functional Ste20 kinase. Its involvement in the Hippo signalling pathway was further verified through interaction and phosphorylation assays with the two newly identified FvMOB1A and FvMOB1B, homologs of the core component MOB1 in this pathway. Additionally, FvM4K1-RNAi knockdown and overexpression (OE) lines were generated and analyzed, revealing that FvM4K1 positively regulates organ and fruit size in woodland strawberry.

## Results

### FvM4K1 Contains the Conserved Ste20 Kinase Domain

The genome of woodland strawberry (*F. vesca*, ‘Hawaii 4’) was searched using BLAST with the amino acid sequence of the kinase domain of Arabidopsis AtSIK1 (At1g69220), a Ste20 homolog (Xiong et al., 2016), resulting the identification of FvM4K1 (FvH4_4g29800). A maximum likelihood (ML) phylogenetic tree was subsequently constructed using the amino acid sequences of the kinase domains of FvM4K1 and other known Ste20p homologs, including Cdc15p (YAR019C, P27636) in yeast (*S.cerevisiae*), Hpo (Hippo, Q8T0S6) in fruit fly (*D. melanogaster*), hMst1 and hMst2 (Mammalian Ste20-like kinase 1/2, Q13043, Q13188) in humans, and AtSIK1 in Arabidopsis. FvM4K1 is in the same branch as AtSIK1 and both clustered with Hpo, Mst1 and Mst2 while Cdc15p and Ste20p from yeast are distantly related to FvM4K1 (Fig. 1a).

**Figure 1.**
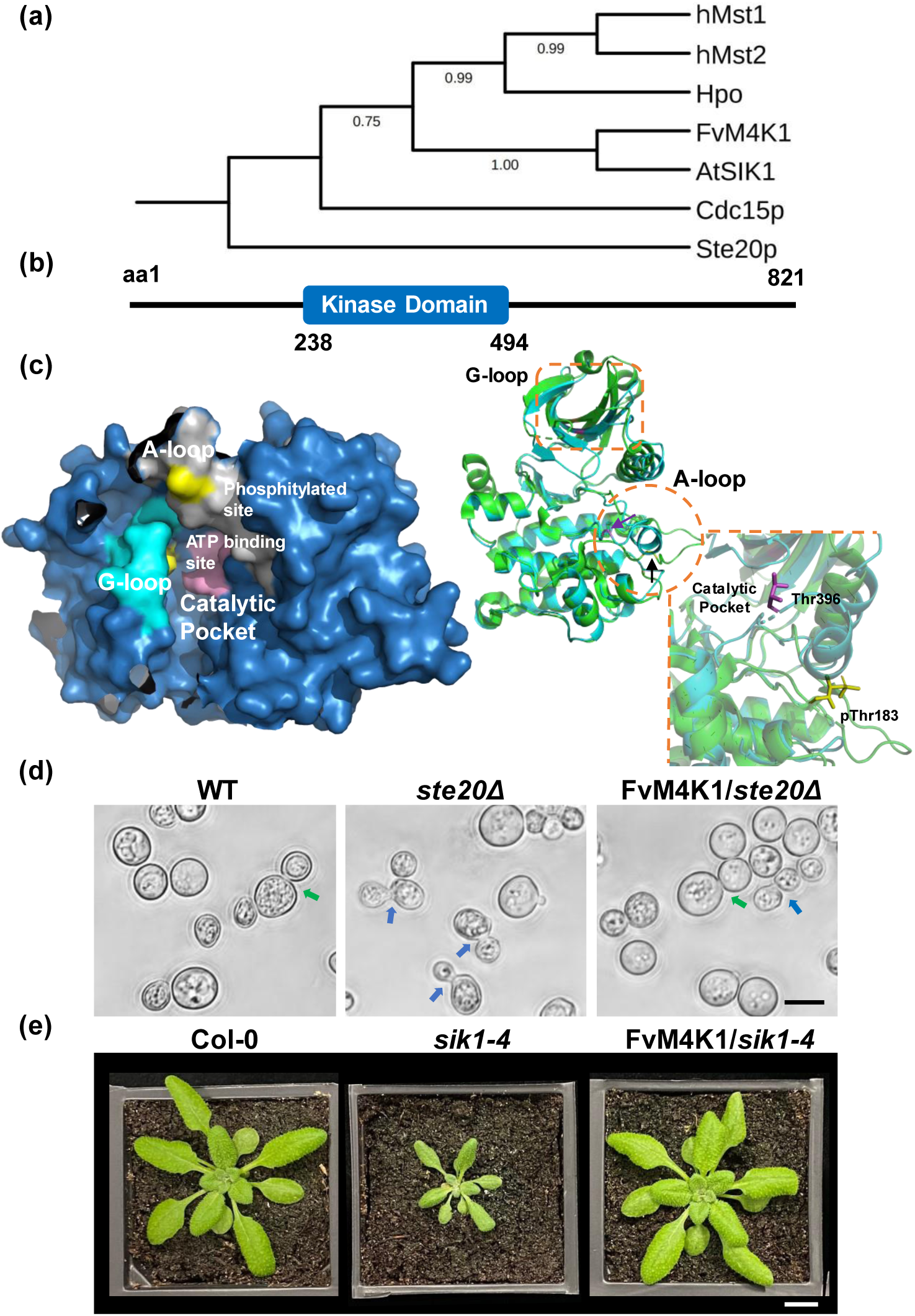
Identification and functional verification of FvM4K1 from woodland strawberry (Fragaria vesca). (a) Phylogenetic tree created by MEGA7.0 from sequence alignment of kinase domains of FvM4K1 of woodland strawberry, AtSIK1 of Arabidopsis, Ste20p and Cdc15p of yeast, Hpo of Drosophila and hMst1/2 of humans. (b) Schematic structure of FvM4K1. The kinase domain between aa238-494 is in blue box. Numbers indicate amino acid positions. (c) Three-dimensional protein structure prediction by AlphaFold. Left panel, the kinase domain of FvM4K1. The G-loop (cyan) and the conserved residue Lys269 for kinase activity close to the G-loop (yellow), the A-loop (grey) and Thr396 (yellow), the auto-p site in the A-loop, the catalytic pocket in the central core and the ATP-binding site (lilac) within it are indicated. Right panel, alignment between the kinase domains of FvM4K1 and phosphorylated Mst1 (3COM in PDB). Green, phosphorylated Mst1; cyan, non-phosphorylated FvM4K1.The A-loop (orange circle) and G-loop (orange box) are highlighted. Lilac line and purple arrow, Thr396 of FvM4K1, yellow line and black arrow, Thr183 of Mst1. (d) FvM4K1 rescues the budding defect of *ste20Δ* mutant. Yeast cells of WT (BY4741), mutant *ste20Δ* and transgenic *ste20Δ* containing FvM4K1 were observed by phase-contrast microscopy. Arrows indicate the budding sites for WT (green) and mutant (blue). Scale bar = 5 μm. (e) FvM4K1 rescues the growth defect of *atsik1-4*, a AtSIK1 loss-of-function T-DNA insertion mutant. Four-week-old WT, *sik1-4* and transgenic *sik1-4* expressing FvM4K1. Scale bar = 10 mm.

Like Ste20 kinases from yeast and animals FvM4K1 also contains the conserved kinase domains (KDs). However, notably the positions of KDs vary significantly between these kinases (Fig. S1a). While Ste20 has a C-terminal KD, Cdc15p, Hpo and Mst1&2 have N-terminal KDs. In contrast, FvM4K1 (spanning amino acids 238-494, Fig. 1b) and AtSIK1 feature KDs positioned one-third away from the N-terminus and two-thirds away from the C-terminus, suggesting that plant Ste20-like proteins may function differently compared to their yeast and mammalian counterparts (Fig. S1a). In addition, eleven major conserved subdomains (I to XI) were identified (black boxes) in the conserved KDs (Fig. S1b). Notably, the glycine-rich loop (G-loop) in subdomain I, with the consensus sequence ‘GXGXXGXX,’ is highly conserved across species. Lys34, identified as the kinase activity site (Taylor and Kornev, 2011), is located in subdomain II. Subdomain VIII includes the activation loop (A-loop) characterized by the consensus sequence ‘KRNT(V/F)(I/V)GTPyWMAPEv’ (purple box), which is recognized as the Ste20 family signature sequence (Delpire, 2009). Within this domain, Thr161 (red box) is a conserved autophosphorylation site (auto p) linked to catalytic activity (Hanks et al., 1988).

AlphaFold 2 prediction of the three-dimensional structure of the kinase domain of FvM4K1 shows that it contains a typical G-loop (cyan) and an A-loop (grey). The conserved residues Lys269, the kinase activity site, is close to the G-loop while Thr396, the auto-p site is in the A-loop (both highlighted in yellow, left panel, Fig. 1c). The catalytic pocket, located in the central core, contains the ATP-binding site (lilac) (Mu et al., 2022). To explore the function of FvM4K1, we compared its KD structure with that of the activated Mst1 containing the phosphorylated Thr183 (Protein Data Bank [PDB] code: 3COM). Interestingly, both structures are largely overlapped, showing they are highly conserved (right panel, Fig. 1c. Green, phosphorylated Mst1; cyan, non-phosphorylated FvM4K1). The only notable difference lies in the A-loop (orange circle) where the KD of FvM4K1 folds into a helix with Thr396 (lilac line and purple arrow) hidden in the pocket while the phosphorylated Thr183 (yellow line and black arrow) of Mst1 is extended outward and exposed (Shi et al., 2015).

Therefore, analysis of the kinase domain of FvM4K1 predicted it to be a member of the Ste20 family with a conserved functional kinase domain.

### FvM4K1 Functions as Ste20 Family Protein

To determine whether FvM4K1 is a homolog of yeast Ste20p, we transformed the Ste20p lacking mutant *ste20Δ* with FvM4K1. As shown in Fig. 1d, the mutant exhibited defects in budding site selection (blue arrows) compared to the wild type (WT) cells (green arrow) (Sheu et al., 2000). However, when FvM4K1 was present approximately 70% of the transgenic cells (FvM4K1/*ste20Δ*) displayed a normal polar budding pattern similar to WT (green arrow). This partial rescue of the budding defect phenotype in *ste20Δ* suggests that FvM4K1 is functionally homologous to Ste20p in yeast.

We then investigated whether FvM4K1 functions similarly to AtSIK1, a Ste20 homolog in Arabidopsis (Xiong et al., 2016). To test this, we transformed the Arabidopsis *atsik1-4* mutant, which lacks *AtSIK1*, with *FvM4K1*. As shown in Fig. 1e and Fig. S2, the *atsik1-4* mutant displayed a dwarfed phenotype, with smaller leaves and shorter roots compared to WT (Xiong et al., 2016). However, *atsik1-4* plants expressing *FvM4K1* were indistinguishable from WT at both the seedling and mature stages, indicating that FvM4K1 fully rescues the growth defects of *atsik1-4* and thus functions like AtSIK1.

### M4K1-RNAi Knock-Down Strawberry Plants Showed Smaller Vegetative and Reproductive Organs

To investigate the biological function of FvM4K1 in woodland strawberry we generated eight lines of M4K1-RNAi transgenic strawberry plants (Fig. S3a). Down-regulation of *FvM4K1* was confirmed by RT-qPCR showing up to 0.8-fold transcript reduction in six of the eight RNAi lines compared to controls (Fig. 2a & S3b). We selected line 5 for further phenotypic analysis due to its lowest transcription of *FvM4K1*. The M4K1-RNAi plants were significantly shorter with smaller leaves and shorter roots compared to WT (Fig. 2c, e-g). Their floral organs, including petals, calyxes, stamens and pistils were also significantly smaller compared to control plants. Specifically, the petal area and pistil length were reduced by ∼23%, and calyx nearly 50% (Fig. 3a, f-i & S3c). The fruits (receptacles) during different developmental stages were also observed given their importance in marketing value. While no differences were observed between RNAi and WT plants at 4-8 day-after-flowering (DAF) significant differences appeared from 14-16 DAF, with the most notable size reduction at 22-24 DAF where RNAi fruits were only half the size of control fruits (Fig. 2e, left panel, j & S3d-e). Additionally, the seeds (achenes) of RNAi fruits were significantly smaller and lighter than controls, although seed length did not show a significant difference (Fig. 2e, right panel, S3f-g).

**Figure 2.**
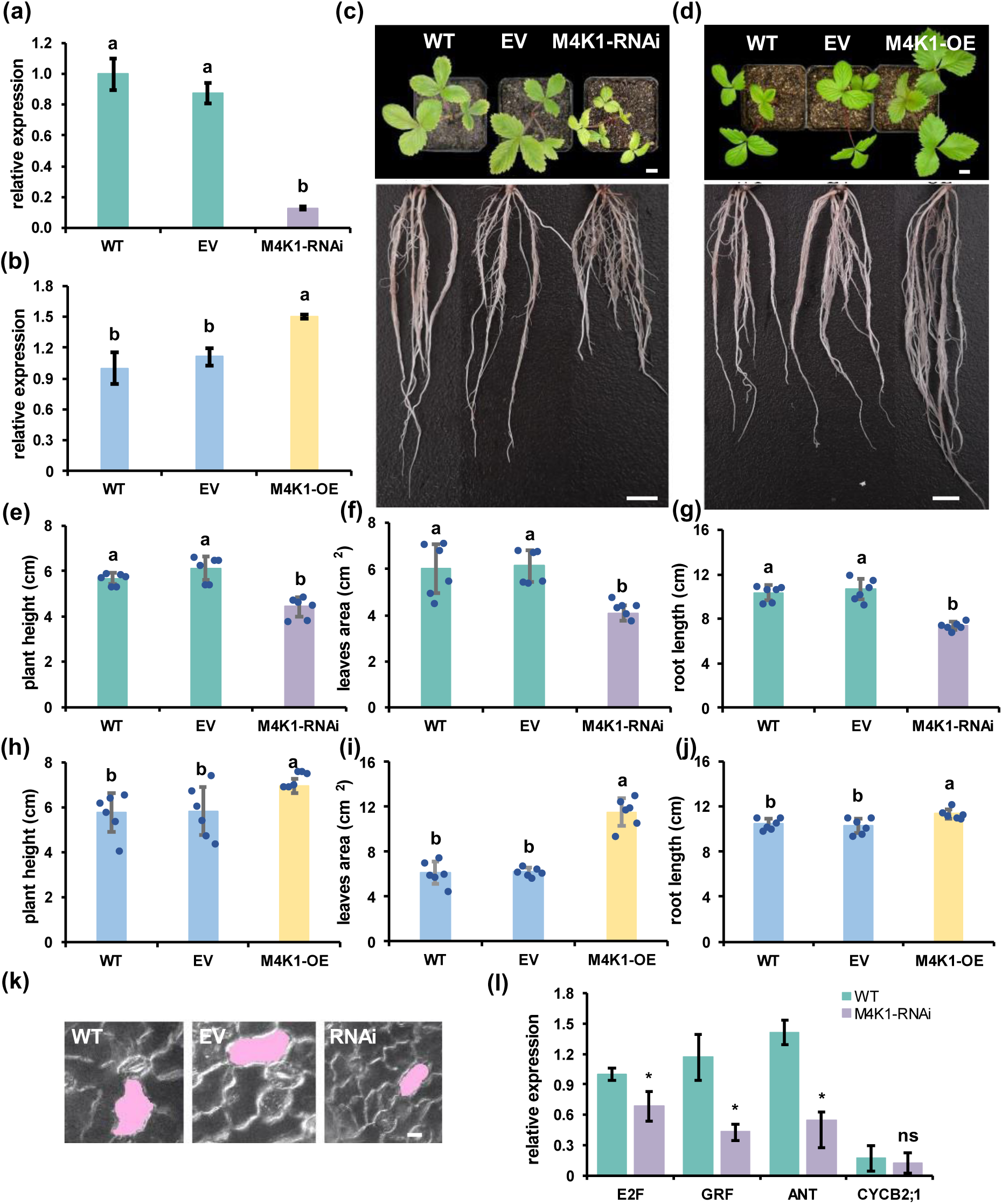
FvM4K1 regulates vegetative organ size in woodland strawberry (*Fragaria vesca*). (a) Detection by RT-qPCR the transcript level of *FvM4K1* in the WT, transgenics harbouring the empty vector (EV) and M4K1-RNAi. (b) Detection by RT-qPCR the transcript level of *FvM4K1* in the WT, transgenics harbouring the empty vector (EV) and 35S:M4K1 OE plants. The relative transcript level of FvM4K1 was calculated by 2^-ΔΔCt^ method using *FvActin* (FvH4_7g22410) as reference gene and transcript level of WT as 1. Error bars = means ± SD (n=3). The different letters indicate significant difference between samples at p < 0.05 calculated by one-way ANOVA with Duncan test using software SPSS. (c) 8-week-old WT, EV and M4K1-RNAi transgenic strawberry plants. Scale bar = 10 mm. (d) 8-week-old WT, EV and 35S:M4K1 OE strawberry plants originated from T0 stolons. Scale bar = 10 mm. (e-g) Measurements of plant height, leaf area and root length of WT, EV and M4K1-RNAi in (c). (h-j) Measurements of plant height, leaf area and root length of WT, EV and M4K1-OE in (d). Error bars are means ± SD (n=6) in (e-j). Different letter indicates statistically significant difference between RNAi, WT and EV plants at p < 0.05 calculated by one-way ANOVA with Duncan test using software SPSS. (k) Lower epidermal cells of leaves. A representative cell from each genotype is shaded with pink. Scale bar = 10 um. (l) Detection by RT-qPCR the transcript level of core cell cycle marker genes and regulators in 14-day-old seedlings of WT and transgenic M4K1-RNAi plants. E2FD, G1/S specific marker; CYCA2;1, S/G2 specific marker; GRF and ANT, transcription factors that regulate cell size and cell number. The relative transcript level was calculated by 2^-ΔΔCt^ method using *FvActin* (FvH4_7g22410) as reference gene and transcript level of *E2FD* in WT as 1. Error bars = means ± SD (n=3). * indicates difference between samples at p< 0.05 in t-test; ns, no significant difference.

**Figure 3.**
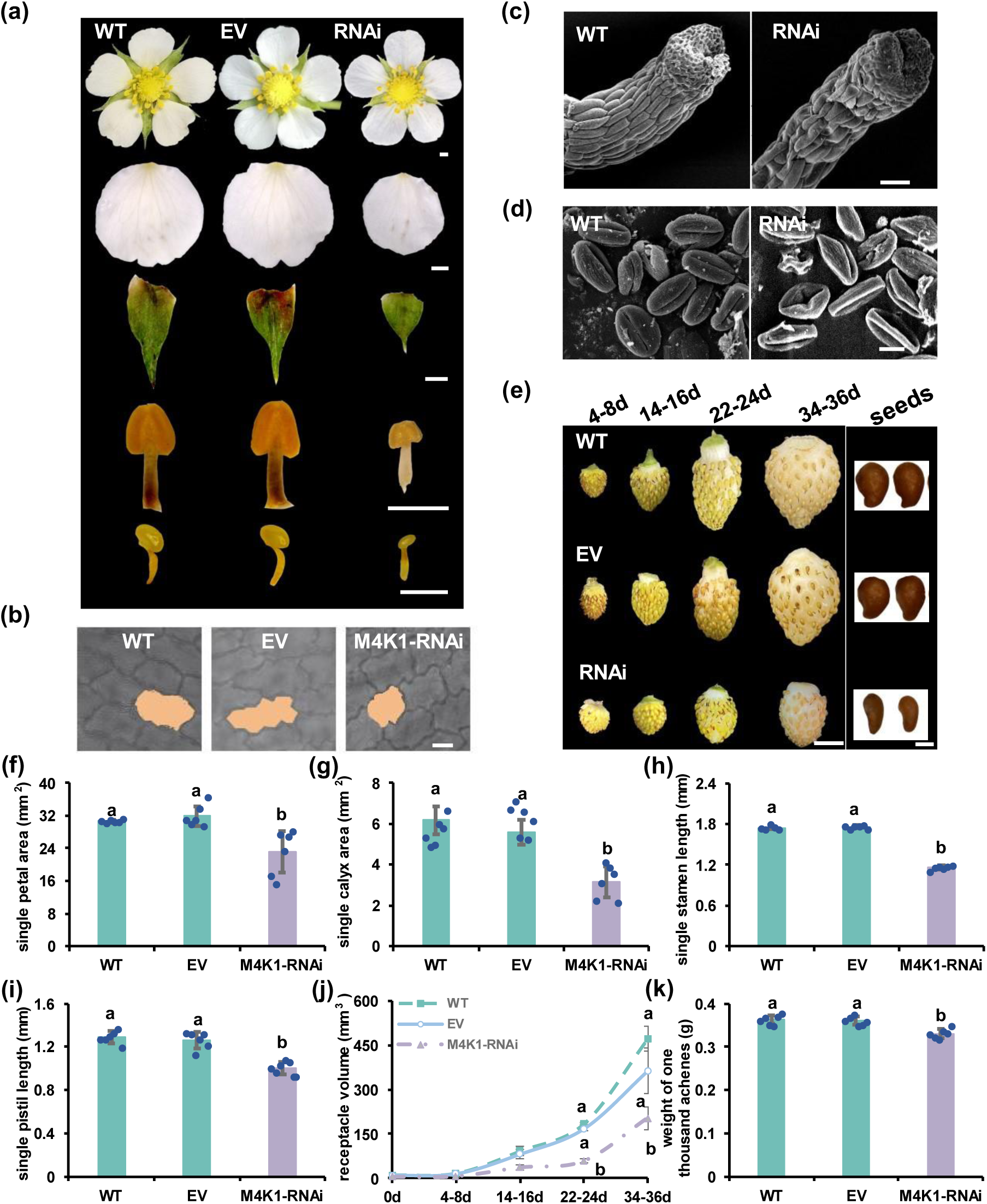
FvM4K1 regulates reproductive organ size of woodland strawberry (*Fragaria vesca*). (a) Fully open flowers, petals, calyx, stamen and pistil from WT, EV and M4K1-RNAi. Scale bar = 1 mm. (b) The epidermal cells of petals from WT, EV and M4K1_RNAi plants. A representative cell from each genotype is shaded with orange. Bar = 10 µm. (c) Scanning electron microscope (SEM) observation of styles. Scale bar = 50 um. (d) Pollen grains by SEM. Scale bar = 10 um. (e) Fruits (receptacles) at different developmental stages (days after flowering) and seeds (achenes). Scale bar = 5 mm in the left panel and = 0.5 mm in the right panel. (f-i) Measurements of petal area, calyx area, stamen length and pistil length. (j-k) Measurements of the volume of the receptacles (fruits) at different developmental stages and weight of thousand achenes in mature achenes (seeds). Error bars are means ± SD (n=6). Different letter indicates statistically significant difference between RNAi, WT and EV plants at p < 0.05 calculated by one-way ANOVA with Duncan test using software SPSS.

These results clearly demonstrate that reducing *FvM4K1* expression via RNAi leads to reduced size in both vegetative and reproductive organs in strawberry plants.

### FvM4K1-RNAi Plants Had Reduced Cell Size and Numbers Compared to WT

We measured the lower epidermal cells of FvM4K1-RNAi leaves, revealing that they were significantly smaller (630 ± 73 μm², n=10) than those of WT (975 ± 101 μm², n=10) (Fig. 2k & S5a). Additionally, the number of pavement cells in M4K1-RNAi leaves was 629,600 ± 13,026 cells (n=6) compared to 696,966 ± 28,890 in WT (n=6), indicating that the RNAi plants had fewer cells than WT (Fig. S5b). Consistent with these results, the cell size and number of RNAi petals were also significantly reduced (Fig. S5c,d).

We further examined the transcript levels of genes associated with the cell cycle (*E2F* and *CYCB2;1*) and the key transcription factors *Growth-Regulating Factor* (*GRF*) and *AINTEGUMENTA* (*ANT*) in 14-day-old seedlings of WT and RNAi plants. As shown in Fig. 2l, although the transcript level of *CYCN2;1* was not significantly different, *E2F*, a marker for G1/S and G2/M cell cycle transitions (Desvoyes and Gutierrez, 2020) was reduced in RNAi plants compared to WT. Notably, the transcripts of *GRF* and *ANT*, which are positive regulators of cell size and cell number (Mizukami and Fischer, 2000; Omidbakhshfard et al., 2015), were downregulated by 0.6-fold in RNAi compared to control plants. Thus, the expression of genes involved in both cell cycle and its regulation are reduced in the M4K1-RNAi plants, resulting in reduction in both cell size and cell number.

### Down-regulation of FvM4K1 Results in Abnormal Pollen Grains and Pistils

Scanning electron microscopy (SEM) analysis of pollen grains revealed that while WT pollen displayed a regular, plump structure, the majority of pollen grains from M4K1-RNAi plants appeared shrunken and collapsed (Fig. 2d). Abnormalities were also observed in the RNAi pistils, where the style cells were misaligned and many had collapsed, compared to the smooth, striated, and regularly aligned style cells in WT plants (Fig. 2c). Consistent with this the RNAi strawberry plants produced less seeds and majority of the seeds stopped developing at 15 DAF (Fig S8). These findings suggest that FvM4K1 plays a critical role in the development of pollen and pistils during reproduction in strawberry plants.

### FvM4K1-OE Plants Had Bigger Vegetative Organs with Increased Cell Size and Number

We further generated overexpression transgenic strawberry plants M4K1-OE (Fig. S4a-b & 2b). Line 4 with the highest *FvM4K1* expression was further phenotyped, showing that the plants were much larger with bigger leaves and longer roots compared to controls (Fig. 2b, h-j). The cell size and number of the lower epidermis of FvM4K1-OE leaves were significantly increased (Fig. S5e-g). Interestingly, although the OE plants produced bigger achenes (seeds) (Fig. S4c-d) they failed to germinate (Fig. S4f). Therefore, overexpression of FvM4K1 resulted in overgrowth of cells and lethality in strawberry.

### FvM4K1 Is a Kinase and Is Auto-Phosphorylated at Lys269 and Thr396

Members of the Ste20 family are known to undergo auto-phosphorylation (Glantschnig et al., 2002; Wu et al., 2003). To determine whether FvM4K1 is also auto-phosphorylated and at which sites we performed *in vitro* kinase assay. We expressed a truncated ΔNFvM4K1, lacking the N-terminus, because the full-length FvM4K1 could not be expressed in the *E. coli* strain BL21(DE3) (Fig. S6a). Additionally, we expressed a kinase-dead mutant, ΔNFvM4K1^K269E^ as Lys269 is the predicted conserved ATP-binding site (Fig. 1c). The anti-phospho-threonine (anti-pT) antibody detected phosphorylation in ΔNFvM4K1, but not in ΔNFvM4K1^K269E^ (Fig. 4a). Thr396 of FvM4K1 within the A-loop is conserved in Hpo of Drosophila and Mst1/2 of mammals and this site is the auto-phosphorylation site (Fig. 1c) (Deng et al., 2013; Ni et al., 2013; Praskova et al., 2004). Therefore, we mutated Thr396 to Ala (T396A) and generated ΔNFvM4K1^T396A^. We also combined both mutations and generated double mutant ΔNFvM4K1^K278ET396A^. The *in vitro* kinase assay showed that ΔNFvM4K1^T396A^ displayed very weak auto-phosphorylation activity (weak band, Fig. 4a) while no phosphorylation was detected in the double mutant ΔNFvM4K1^K269ET396A^ (no band, Fig. 4a). These results clearly demonstrated that FvM4K1 a threonine kinase, both Thr396 and Lys269 play essential roles in phosphorylation of FvM4K1.

**Figure 4.**
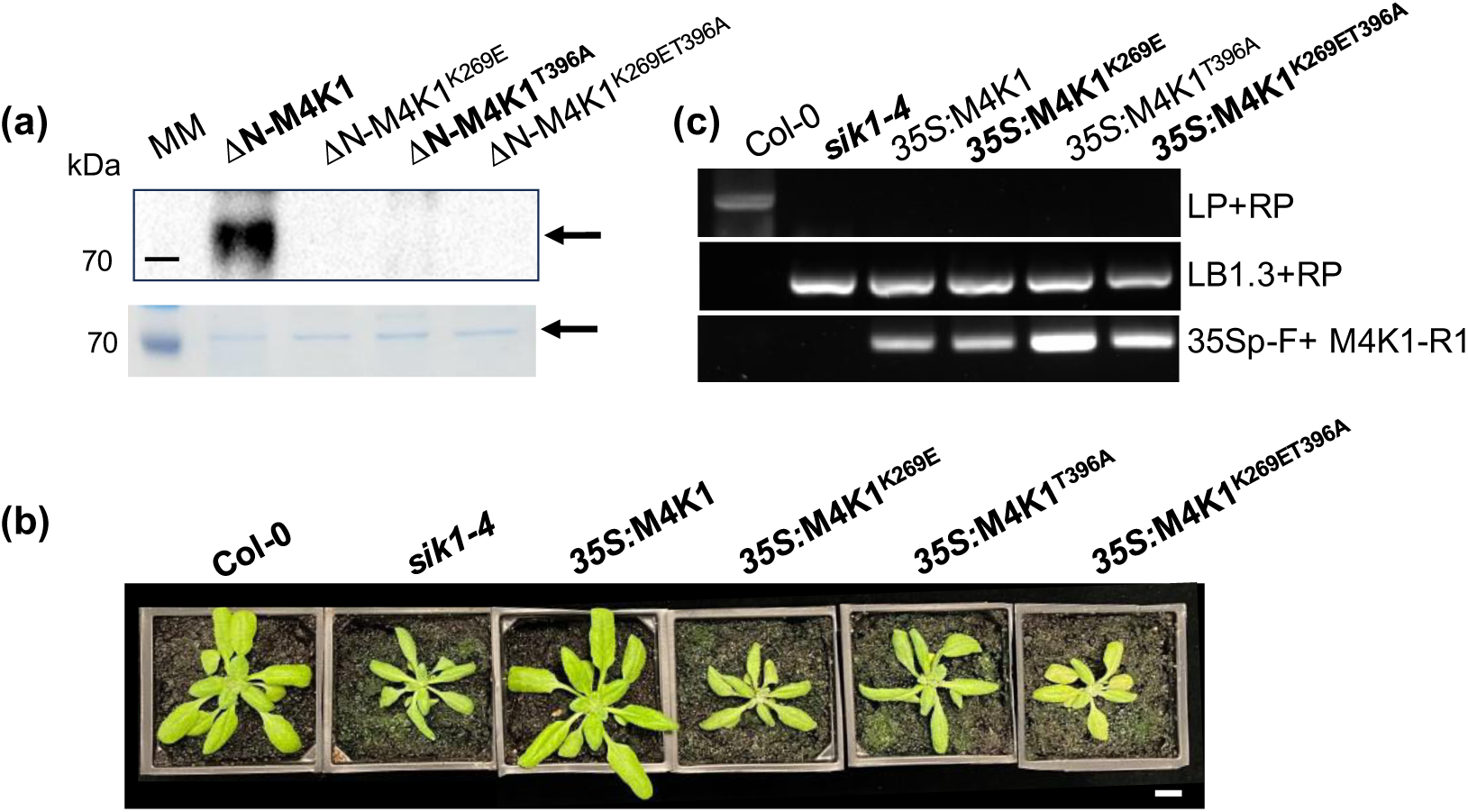
FvM4K1 is auto-phosphorylated and its kinase activity relies on K269 and T396 for function. (a) In vitro kinase assay. Top panel, Western blot detection with anti-phosphor-threonine antibody. A strong band for ΔN-FvM4K1 was detected while no band detected in lanes containing the kinase-dead mutant ΔN-FvM4K1^K278E^, ΔNFvM4K1^T396A^ and the double mutant ΔN-FvM4K1^K269ET396A^. Bottom panel, SDS-PAGE gel stained with EZBlue, showing equal amounts of proteins loaded for ΔN-FvM4K1 and its pointed mutants. Proteins were expressed in BL21(DB3) and purified. Arrows indicate the positions of FvM4K1 and its variants. MM, protein molecular marker. (b) Four-week-old Arabidopsis plants of Col-0, *sik1-4* and transgenic *sik1-4* expressing FvM4K1, FvM4K1^K269E^, FvM4K1^T396A^ and FvM4K1^K269ET396A^. Scale bar = 10 mm. (c) Genotyping by PCR to confirm *sik1-4* background and the presence of *FvM4K1* in the transgenic plants. Primers sik1-4-LP, sik1-4-RP, LBb1.3, 35Sp-F and M4K1-R1 are listed in Table S1.

### Kinase Activity of FvM4K1 is Essential for Organ Size Control

To validate the *in vitro* kinase assay results we transformed FvM4K1 and its single and double mutants FvM4K1^K269E^, FvM4K1^T396A^ and FvM4K1^K269ET396A^ in the Arabidopsis *atsik1-4* mutant. While FvM4K1 fully rescued the growth defects of *atsik1-4*, mutants expressing FvM4K1^K269E^ and FvM4K1^K269ET396A^ were indistinguishable from the *atsik1-4* mutant, showing no rescue of growth (Fig. 4b and S6b-d). Interestingly, FvM4K1^T396A^ provided partial rescue during early stages of growth as indicated by the larger rosette diameter at 4-week-old plants. However, the effect was diminished in older plants, and the 9-week-old plants exhibited similar phenotype as *atsik1-4*, indicating a loss of rescue over time (Fig. 4b & S6b-d).

Therefore, FvM4K1 is a functional kinase, its kinase activity and auto-phosphorylation rely on Lys269 and Thr396, respectively. Importantly, both kinase activity and auto-phosphorylation are essential for the role of FvM4K1 in regulating organ size.

### FvM4K1 Interacts with FvMOB1A and FvMOB1B, the Core Component of the Hippo Pathway

In addition to the Ste20 kinase, Mats/Mob1 has been identified as a key component of the Hippo pathway to regulate organ size in both flies and mammals (Praskova et al., 2008; Wu et al., 2003). Homologs of Mob1/Mats have also been found in Arabidopsis, i.e., AtMOB1A (AT5G45550) and AtMOB1B (AT4G19045) (Xiong et al., 2016). Using the protein sequences of AtMOB1A&B to search the strawberry genome we identified two homologues, FvMOB1A (FvH4_5g23830) and FvMOB1B (FvH4_1g08580) (Fig. S7b). Transcription of *FvMOB1A* was reduced by 60% while *FvMOB1B* was undetectable in the FvM4K1-RNAi plants (line 5), suggesting that both FvMOB1A and FvMOB1B function in an FvM4K1-dependent manner.

To see if FvM4K1 interacts with FvMOB1A&B we conducted a yeast two-hybrid (Y2H) assay. The suitability of Y2H was confirmed by self-activation test as no colonies grew on SD/−T/−L/−A/AbA media (Fig. S7a). In the experimental assay, colonies harboring protein pairs between FvM4K1 and FvMOB1s grew well on nutrient-deficient SD/−T/−L media, indicating gene expression (Fig. 5b & S7c). Colonies expressing FvM4K1-FL/FvMOB1A, FvM4K1-FL/FvMOB1B, and FvM4K1-N/FvMOB1s on selective media (SD/−T/−L/−H/−A/AbA/X-α-Gal) grew well and turned blue, while those containing FvM4K1-N1, -N2, or -C with FvMOB1s struggled to grow and did not turn blue regardless whether the proteins were fused to AD or BD (Fig. 5b & S7c). This indicates that FvM4K1 interacts with FvMOB1A and FvMOB1B, and this interaction depends on an intact kinase domain within the N-terminus of FvM4K1.

**Figure 5.**
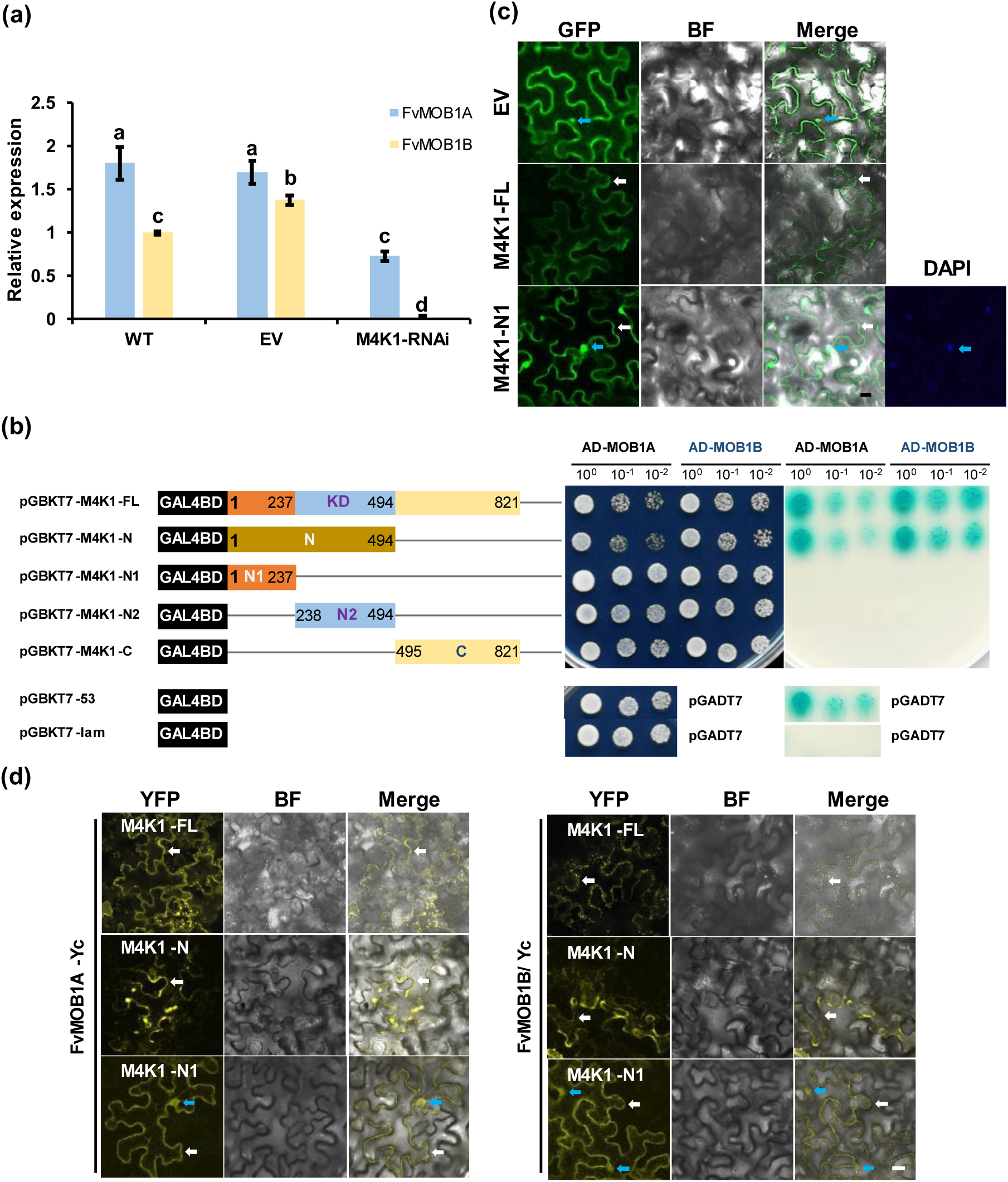
FvM4K1 interacts with FvMOB1A and FvMOB1B. (a) The transcript of *FvMOB1A* and *FvMOB1B* are downregulated in the M4K1-RNAi strawberry plants. The relative transcript levels were calculated by the 2^-ΔΔCt^ method using *FvActin* (FvH4_7g22410) as the reference gene and expression level of *FvMOB1B* in WT as 1. Error bars = means ± SD (n=3). The different letters indicate significant difference between samples at p < 0.05 calculated by one-way ANOVA with Duncan test using SPSS. (b) Y2H. Left panel, schematic structure of the FvM4K1 gene and fragments; Middle panel, colonies grew on the selective media lacking Trp and Leu (SD/−T/−L); right panel, colonies grew on media lacking Trp, Leu, His and Ala and supplemented with 200 ng/mL of Aureobasidin A (AbA) and 40 µg/mL of X-α-Gal (SD/−T/−L/−H/−A/AbA/X-α-Gal). All colonies were grown to reach OD600=1, dilutions of 1/10, 1/100 were spotted on the plates. pGBKT7-53 and pGADT7-T were positive control pair while pGBKT7-lam pGADT7-T negative control. KD, kinase domain. (c) Subcellular localization of FvM4K1-FL and –N1. FvM4K1-FL localization was detected on the PM while FvM4K1-N1 was both on the PM and in the nucleus. Blue arrows indicate nucleus, white arrows indicate PM. Scale bar = 20 µm. (d) BiFC. Top panel, FvMOB1A-YFPc and FvM4K1- and its fragments-YFPn; Bottom panel, FvMOB1B-YFPc and FvM4K1- and its fragments-YFPn. Blue arrows indicate nucleus, white arrows indicate PM. Scale bar = 20 µm.

### Interactions between FvM4K1 and FvMOB1s Occur at Both the Plasma Membrane and Nucleus Which Require an Intact FvM4K1 N-terminus

The interaction between FvM4K1 and FvMOB1 proteins was further confirmed in tobacco leaves using bimolecular fluorescence complementation (BiFC). As shown in Fig. 5d and S7d, yellow fluorescent protein (YFP) signals were detected at the plasma membrane (PM; white arrows) when FvM4K1-FL was co-expressed with either FvMOB1A or FvMOB1B, as well as when FvM4K1-N was co-expressed with FvMOB1A or FvMOB1B, indicating interactions at the PM. Notably, YFP signals were also observed at both the PM and nucleus (blue arrows) when FvM4K1-N1 was co-expressed with FvMOB1A or FvMOB1B, suggesting that nuclear localization of FvM4K1 occurs independent of its kinase domain.

We further assessed the subcellular localization of FvM4K1 and FvM4K1-N1 by expressing their GFP fusions in tobacco leaves. Confocal microscopy revealed that FvM4K1 localizes to the PM (Fig. 5c), while FvM4K1-N1-GFP was observed both at the PM (white arrows) and in the nucleus (blue arrows, confirmed by DAPI staining). These results suggest that FvM4K1 localizes to the PM, which is crucial for its interaction with FvMOB1 proteins.

### FvM4K1 Phosphorylates FvMOB1A and FvMOB1B

Since Ste20 family kinases, like Hpo and Mst1/2, are known to phosphorylate their interacting partners, it is likely that FvM4K1 also phosphorylates FvMOB1A and FvMOB1B. To test this, we conducted Y2H and in vitro kinase assays. After ruling out false positives via the Y2H self-activation test (Fig. S7a), we observed colony growth on nutrient-deficient SD/−T/−L media (Fig. 6a, right panel). However, on selective media (SD/−T/−L/−H/−A/AbA/X-α-Gal), colonies expressing combinations of FvM4K1^T396A^/FvMOB1A and FvM4K1^T396A^/FvMOB1B grew well and turned blue, similar to pairs between FvM4K1-FL and FvMOB1s. However, colonies expressing FvM4K1^K269ET396A^/FvMOB1A and FvM4K1^K269ET396A^/FvMOB1B did not grow or turn blue (Fig. 6a, left panel). These findings indicate that mutation in FvM4K1^K269E^ disrupts interaction with FvMOB1s, while FvM4K1^T396A^ retains its interaction capability. Thus, the interaction between FvM4K1 and FvMOB1s depends on Lys269, linked to kinase activity, rather than Thr396 in the A-loop for auto-phosphorylation.

**Figure 6.**
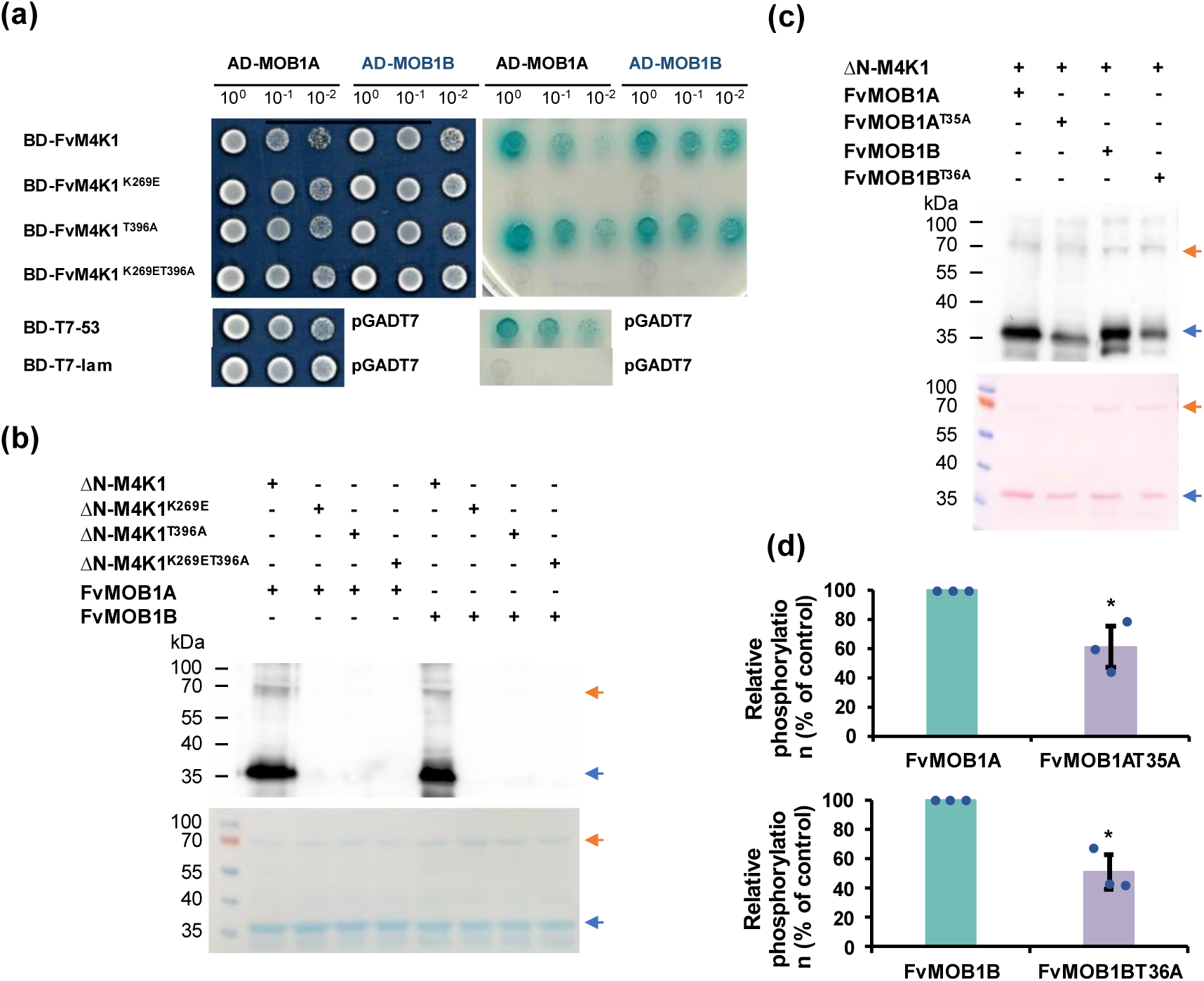
Interaction between FvM4K1 and FvMOB1A&B relies on the kinase activity of FvM4K1. (a) Yeast two hybrid assay. Left panel, yeast colonies on SD/−T/−L selective media showing yeast growth; right panel, yeast colonies on SD/−T/−L/−H/−A/AbA/X-α-Gal showing interaction. Aureobasidin A (AbA, 200 ng/mL) and X-α-Gal (40 µg/mL) were added to the media. All colonies were grown to reach OD600=1, dilutions of 1/10, 1/100 were spotted on the plates. pGBKT7-53 and pGADT7-T were positive control while pGBKT7-lam pGADT7-T negative control pair. (b) FvMOB1A and FvMOB1B are phosphorylated by FvM4K1 detected by *In vitro* kinase assay. Top panel, detection by Western blot with anti-phosphor-threonine antibody; lower panel, EZBlue stained gel. Orange arrows indicate the positions of ΔNFvM4K1 and blue arrows FvMOB1A and FvMOB1B. All proteins were expressed in BL21(DB3) and purified. (c) FvMOB1A and FvMOB1B are phosphorylated at Thr35 and Thr36, respectively, determined by i*n vitro* kinase assay. Top panel, detection by Western blot with anti-phosphor-threonine antibody; bottom panel, Ponceau S stained blot. Orange arrows indicate the positions of ΔNFvM4K1 and blue arrows FvMOB1A and FvMOB1B. All proteins were expressed in BL21(DB3) and purified. (d) Quantification of phosphorylation of WT and mutants of FvMOB1A and FvMOB1B in (C). The phosphorylation (band intensity) of FvMOB1A and FvMOB1B on the blot was regarded as 100%. The phosphorylation of mutant AtMOB1s^TA^ was calculated as the percentage of the peak intensity of AtMOB1s. Error bars are means ± SD (n=3). * indicates difference between samples at p< 0.05 in t-test.

To directly demonstrate role of phosphorylation in the interaction, we expressed and purified the relevant proteins and conducted an in vitro kinase assay. As shown in Fig. 6b, both FvMOB1A and FvMOB1B were detected with an anti-phospho-threonine antibody (anti-pT) (arrows in the first and fifth lanes) when ΔNFvM4K1 was present, confirming that FvM4K1 phosphorylates both FvMOB1A and FvMOB1B (top panel). In contrast, only faint or no bands were observed for FvMOB1A and FvMOB1B when FvM4K1^T396A^ or FvM4K1^K269E^ were used, indicating these mutants had little or no ability to phosphorylate FvMOB1s.

These findings demonstrate that the interaction between FvM4K1 and FvMOB1s depends on both trans- and auto-phosphorylation of FvM4K1.

### Identification of Phosphorylation Sites in FvMOB1A and FvMOB1B by FvM4K1

To identify the specific amino acid residues in FvMOB1A and FvMOB1B were phosphorylated by FvM4K1, we looked into previous studies on hMob1, which identified Thr12 and Thr35 as phosphorylation sites, with Thr35 being more impactful (Praskova et al., 2008). Notably, Thr35 is conserved in FvMOB1A and as Thr36 in FvMOB1B (Fig. S7b). We therefore mutated these residues to alanine (FvMOB1A^T35A^ and FvMOB1B^T36A^) and expressed the mutant proteins in *E. coli* (Fig. 6c, bottom panel). After the kinase reactions where 200 ng of FvM4K1 and 1 µg of FvMOB1A, FvMOB1A^T35A^, FvMOB1B, or FvMOB1B^T36A^, were added all four FvMOB1s could be detected by Western blot with anti-pT antibody. However, phosphorylation of FvMOB1A^T35A^ and FvMOB1B^T36A^ was reduced by 40% and 50% compared to their wild-type counterparts (Fig. 6c, top panel, 6d). Thus, FvM4K1 phosphorylates both FvMOB1A and FvMOB1B, with Thr35 in FvMOB1A and Thr36 in FvMOB1B as key phosphorylation sites.

## Discussion

### FvM4K1 Functions as a Typical Ste20-like Protein in Woodland Strawberry

We identified FvM4K1 (FvH4_4g29800), a Ste20-like protein, in woodland strawberry (*Fragaria vesca*, ‘Hawaii 4’) (Fig. 1). FvM4K1 rescued the growth defect of a yeast *ste20Δ* mutant lacking Ste20p (Fig. 1d), similar to how Arabidopsis AtSIK1 can restore the same mutant (Xiong et al., 2016). This suggests that plant Ste20-like proteins, such as FvM4K1 and AtSIK1, retain the conserved function of Ste20p in regulating organ size via cell division. The role of Ste20 family kinases in controlling organ size through the Hippo pathway is well-established in flies and mammals (Lu et al., 2010a; Wu et al., 2003). In humans, Ste20-like proteins Mst1/2 auto-phosphorylate and trans-phosphorylate substrates at serine and threonine residues (Creasy and Chernoff, 1995a; Creasy and Chernoff, 1995b). While AtSIK1, a Ste20 homolog in Arabidopsis, has been shown to autophosphorylate, its ability to trans-phosphorylate substrates has not been reported (Zhang et al., 2018).

To determine whether FvM4K1 undergoes autophosphorylation, we performed *in vitro* kinase assays. Detection with an anti-phospho-threonine antibody confirmed FvM4K1 autophosphorylation, establishing it as a threonine kinase (Fig. 4a). Lys269 within the conserved ATP-binding site is conserved with other Ste20 kinases (Fig. 1c). Phosphorylation is absent in kinase-dead mutants such as Hpo^K71R^ in Drosophila and Mst1^K59R^ in mice (Glantschnig et al., 2002; Wu et al., 2003), underscoring the importance of this lysine residue in kinase activity. Consistent with this the ability of FvM4K1^K269E^ to auto-phosphorylate and trans-phosphorylate FvMOB1A and FvMOB1B were abolished (Fig. 4a & 6b-c), confirming FvM4K1 as a threonine kinase. Thr396 in the A-loop of FvM4K1 (Fig. 1c & S1) is also conserved with Thr183/Thr180 in Mst1/2 and Thr195 in Hpo (Glantschnig et al., 2002; Jin et al., 2012). Mutations at these sites in Mst1/2 significantly reduce phosphorylation (Glantschnig et al., 2002; Ni et al., 2013; Praskova et al., 2008). Similarly, we found that both autophosphorylation and trans-phosphorylation activities were abolished in the FvM4K1^T396A^ mutant (Fig. 4a & 6b-c), demonstrating that auto-phosphorylation at Thr396 is essential for FvM4K1 function.

Thus, the lysine residue in subdomain II (Lys269) and the threonine residue in the A-loop (Thr396) are crucial for the kinase activity and autophosphorylation of FvM4K1. These structural features are conserved across Ste20 kinases in plants, yeast, flies, and mammals although their positions are different due to the positions of the kinase domains in these proteins (Fig. 1c & S1).

### The Role of the Hippo Pathway in Strawberry: Kinase-Scaffold Interactions and Phosphorylation Dynamics in Plants

The Hippo pathway is a critical signalling mechanism that regulates organ size by controlling kinase-scaffold protein interactions and phosphorylation events. A Ste20 kinase interacts with a WW-domain scaffold protein, leading to phosphorylation of downstream complexes, including Mob1 family proteins and LAT/NDR kinases, which in turn regulate target proteins involved in organ size control (Zhao et al., 2011; Zheng and Pan, 2019). While well-characterized in Drosophila and mammals, the core components of the Hippo pathway in plants are less understood. In Arabidopsis, three components have been identified, AtSIK1 (Ste20 homolog), AtMOB1A/B (Mob1 homologs) and eight AtNDRs (LAT/NDR homologs) (Guo et al., 2020; Xiong et al., 2016; Yoon et al., 2021). Both AtMOB1A and AtMOB1B interact with AtSIK1 and NDR2/4/5, similar to the conserved interactions found in animals (Praskova et al., 2008; Wei et al., 2007). This suggests that the Hippo pathway shares conserved components and interactions between plants and animals.

Although the AtSIK1-AtMOB1A/B interactions have been confirmed in Arabidopsis, the nature of this interaction remains unknown (Xiong et al., 2016). To address this, we performed yeast two-hybrid assays and demonstrated that their homologs, FvM4K1 and FvMOB1A/B in woodland strawberry also interact. This interaction depends on the intact N-terminus of FvM4K1, as neither the N1 fragment (lacking the kinase domain) nor the N2 fragment (containing only the kinase domain) interacted with FvMOB1s (Fig. 5b & S7c). We further validated the interactions using bimolecular fluorescence complementation assays in tobacco, although a discrepancy emerged showing an unexpected interaction between FvM4K1-N1 and FvMOB1s (Fig. 5d). Importantly, the interaction between FvM4K1 and FvMOB1s is kinase-dependent, as no interaction was observed between the kinase-dead mutant FvM4K1^K269E^ and FvMOB1s in Y2H assays (Fig. 6a). This was further corroborated by *in vitro* kinase assays where FvMOB1A and FvMOB1B were phosphorylated by FvM4K1. A mutation at Thr396 in FvM4K1 significantly reduced its ability to phosphorylate FvMOB1s (Fig. 6b), indicating that the kinase activity of FvM4K1 is self-regulated through autophosphorylation.

In humans, Mst1 phosphorylates hMob1A at two key sites, Thr12 and Thr35, with Thr35 having a more profound effect on Mob1A phosphorylation (Praskova et al., 2004). Similarly, phosphorylation of FvMOB1A^T35A^ and FvMOB1B^T36A^ was reduced by up to 50% (Fig. 6c & d), suggesting that these threonine residues are critical phosphorylation sites. Interestingly, the auto-phosphorylated Mst2 binds hMob1A and its mutant hMob1A^T12A/T35A^ more effectively than inactive Mst2. However, in the presence of ATP and Mg²⁺, Mst2 binding to hMob1A decreased significantly, while the binding to hMob1A^T12A/T35A^ remained unaffected^23^. This suggests that phosphorylation of hMob1A by Mst2 promotes complex dissociation. A similar dynamic may occur between FvM4K1 and FvMOB1s, where phosphorylation is critical for interaction, but other structural elements likely contribute to the stability of the interaction complex. This, of course, remains to be determined.

### Divergent Roles of the Hippo Pathway in Organ Size Regulation: FvM4K1-Mediated Growth in Strawberry and Functional Comparisons with Animal Systems

To investigate the role of FvM4K1 in woodland strawberry, we generated FvM4K1-RNAi knockdown plants. One RNAi line (line 5) with significantly reduced *FvM4K1* expression exhibited reduced stature with smaller vegetative and reproductive organs which resulted from reduction in both cell size and cell number (Figs. 2, 3 & S3, S5). In contrast, plants overexpressing FvM4K1 (M4K1-OE) displayed increased height and larger leaves, associated with increased cell size and number (Figs. 2d, h-j & S4, 5). These results suggest that FvM4K1 regulates organ size in strawberry by promoting cell expansion and proliferation. Similarly, Arabidopsis mutants, such as *atsik1-4*, lacking AtSIK1, show stunted growth with smaller organs caused by reduced cell size and number compared to wild-type plants (Xiong et al., 2016). Furthermore, introducing FvM4K1 into *atsik1-4* mutants rescued their growth defects, but neither the kinase-dead mutant FvM4K1^K269E^ nor the phospho-threonine mutant FvM4K1^T396A^ did (Fig. 1e & S2), indicating that FvM4K1 in strawberry functions similarly to AtSIK1 in Arabidopsis, playing a positive role in cell division and expansion. This contrasts with the role of Ste20 family kinases in animals, where loss-of-function mutations often result in overgrowth. For example, in fruit fly, mutations in Hpo kinase lead to tissue overgrowth, resulting in larger organs such as larger eyes, due to excessive cell proliferation (Wu et al., 2003). Similarly, overexpressing Hpo phosphorylation mutants T195A or kinase-dead K71R causes enlarged wings while wild-type flies exhibit smaller and more compact wings (Jin et al., 2012). Thus, in animals, Ste20 family kinases act as negative regulators of cell proliferation. This indicates a fundamental difference in the function of Ste20 family proteins between plants and animals, with plants using these proteins to promote while animals use them to suppress growth.

Ste20 kinases are key components of the Hippo signaling pathway, which regulates organ size through interactions with and phosphorylation of other pathway components (Zhao et al., 2011; Zheng and Pan, 2019). Consequently, mutations in these components can also result in growth defects similar to those seen in Ste20 mutants. For example, in *Drosophila*, mutations in *wts* and *sav* lead to tissue overgrowth (Justice et al., 1995; Kango-Singh et al., 2002), while co-expression of Hpo and Mats significantly reduces eye size (Lai et al., 2005; Wu et al., 2003). Conversely, in Arabidopsis, the *mob1a–/– mob1b+/–* mutant, which lacks AtMOB1A in both chromosomes and AtMOB1B in one, shows reduced plant height, shorter roots and siliques (Guo et al., 2020). More severe growth defects are observed in the in *sik1–/– mob1a–/–* and *sik1–/– mob1a+/–* double mutants where growth arrest occurs and seed germination is impaired, indicating growth suppression in these mutants (Xiong et al., 2016). Thus, while the Hippo pathway is evolutionarily conserved across eukaryotes, it appears to play opposite roles in regulating cell proliferation and organ size in plants and animals. In plants, the Hippo pathway promotes growth, whereas in animals, it suppresses growth.

However, the role of the Hippo pathway in reproduction appears to be conserved between plants and animals. In FvM4K1-RNAi strawberry plants, some seeds (achenes) failed to develop and aborted (Fig. S8), likely due to abnormal pollen grains or pistils observed by SEM (Fig. 3c &d). This is consistent with the observation in Arabidopsis where the mutant *atsik1-4* had fewer seeds per silique than WT although the cause remains to be explored (Xiong et al., 2016). Interestingly, overexpression of FvM4K1 in woodland strawberry caused lethality, as T1 achenes from M4K1-OE plants failed to germinate (Fig. S4f). In fly, the functional loss of Hpo in eyes leads to pupal lethality in eyeless flies. Additionally, 66% of *hpo* null mutants die, and none survive when a heterozygous *sav* or *wts* mutation is introduced. Co-expression of *Hpo* and *Wts* causes complete lethality (Wu et al., 2003). These findings suggest a conserved role for the Hippo pathway in reproduction across species.

Based on these findings, we propose a model for the Hippo pathway in woodland strawberry. FvM4K1 positively regulates organ size and this relies on its kinase activity. In its active form (“Hippo on”, Fig. 7, left panel), FvM4K1 undergoes auto-phosphorylation and interacts with and phosphorylates FvMOB1A and FvMOB1B. FvM4K1 may also interact with and phosphorylate NDR family proteins, leading to the activation of transcription factors and the expression of genes that promote cell proliferation and organ enlargement. However, when the pathway is inactive (“Hippo off”, Fig. 7, right panel), FvM4K1 is unable to undergo auto-phosphorylation or trans-phosphorylate downstream components, resulting in reduced organ size.

**Figure 7.**
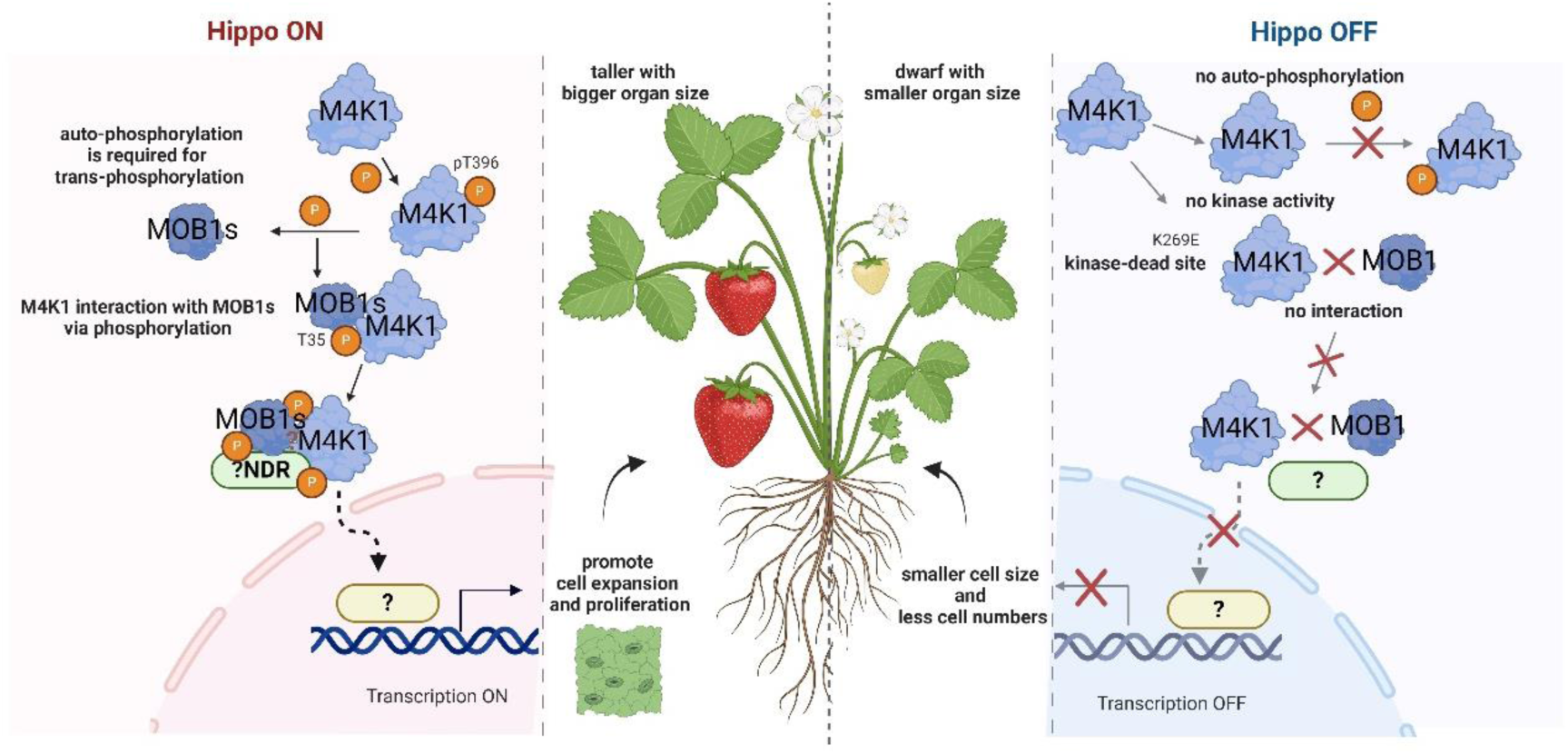
The proposed model of Hippo signalling pathway in organ size control in woodland strawberry (*Fragaria vesca*). The FvM4K1 regulates organ size and this depends on its kinase activity. In its active form (Hippo on, left panel), FvM4K1 interacts with and phosphorylates Hippo pathway components, such as MOB1s, a process facilitated by auto-phosphorylation of FvM4K1 at Thr396. FvM4K1 may also interact with and phosphorylates the NDR family proteins aided by the phosphorylated FvMOB1s. This leads to the activation of transcription factors which enter the nucleus where they can trigger the transcription of genes involved in cell proliferation, resulting in larger strawberry fruits and other organs. However, when this pathway is off (right panel), FvM4K1 is unable to auto-phosphorylate (T396A) and/or does not have kinase activity (K296E, kinase dead). Therefore, interaction and trans-phosphorylation of FvMOB1s and other Hippo pathway components cannot be achieved, genes controlling cell size and number are not activated, thus the observed smaller organ size in strawberry.

In summary, FvM4K1 functions as a kinase in the Hippo signalling pathway in woodland strawberry by interacting with and phosphorylating the scaffold proteins FvMOB1A and FvMOB1B, homologous to those found in animals. Like AtSIK1 in Arabidopsis, FvM4K1 positively regulates organ size in strawberry, contrasting with the typically negative regulatory role of Ste20 family kinases in animals. Although the molecular basis for this functional divergence remains unclear, further investigation into the unique structural chatacteristics of plant Ste20-like proteins and other components of the plant Hippo pathway may shed light on these differences.

## Materials and methods

### Plant Materials

Woodland strawberry (*Fragaria vesca*, ‘Hawaii 4’) was grown under 11h-light (120 μm m^-2^ s^-1^) and 13h-dark at 21°C. Wild-type Arabidopsis (Col-0) and the T-DNA insertion lines *atsik1-4* (SALK_051369) (Xiong et al., 2016) were obtained from the Arabidopsis Biological Resource Centre (ABRC, http://www.arabidopsis.org/abrc/) and grown under a 16h-light (120 μm m^-2^ s^-1^) and 8h-dark cycle at 21°C.

### Bioinformatic Analysis

For phylogenetic analysis, the amino acid sequences of the kinase domain of strawberry FvM4K1 (FvH4_4g29800) was aligned with Ste20p and Cdc15p from yeast (Bardin et al., 2003; Peter et al., 1996), Hpo from fruit fly (Wu et al., 2003), hMst1/2 from humans (Ni et al., 2013; Praskova et al., 2004) and AtSIK1 from Arabidopsis (Xiong et al., 2016) ClustalW in MEGA7.0 software. A maximum likelihood (ML) phylogenetic tree was constructed using the Poisson substitution model and visualized with iTOL (https://itol.embl.de/).

For 3D protein structure prediction, the kinase domain of FvM4K1 was modeled using AlphaFold (https://colab.research.google.com/github/deepmind/alphafold/blob/main/notebooks/AlphaFold.ipynb). The predicted structure with the highest mean pLDDT (88.59) was visualized in PyMOL (Jumper et al., 2021; Varadi et al., 2022), and aligned with the known structure of the Mst1 kinase domain (PDB code: 3COM) to indicate the active conformation of FvM4K1.

### Yeast Complementation Assay

The coding region of *FvM4K1* was cloned into the yeast expression vector pYES-DEST52 using Gateway cloning (Invitrogen) and transformed into wild-type yeast BY4741 and the *ste20Δ* mutant (Y00956) (MATa, ura3Δ0, leu2Δ0, his3Δ1, met15Δ0, YHL007c::kanMX4) (EUROScarf, http://www.euroscarf.de/). Empty vector controls were also transformed into both yeast strains. Colonies were grown on selective media containing 2% galactose to induce protein expression. Cells were observed using phase-contrast microscopy (Zeiss LSM 710).

### Arabidopsis and Strawberry Transformation

C-terminal GFP-tagged FvM4K1 and its mutants (FvM4K1^K269E^, FvM4K1^T396A^, FvM4K1^K269ET396A^) were cloned into pH7FWG2-293 via Gateway cloning (Katzen, 2007; Zhang et al., 2022) and transformed into *atsik1-4* Arabidopsis using the floral dipping method, followed by genotyping (Qi et al., 2004).

For FvM4K1-RNAi strawberry plants, a ∼200 bp fragment from the kinase domain of FvM4K1 was cloned into pK7GWIWG2(II)-RR-277 to generate a hairpin structure (Zhang et al., 2022) and transformed into woodland strawberry. Empty vector controls were also transformed. Overexpression of FvM4K1 was achieved by cloning the full-length gene into pH7FWG2-293 (Zhang et al., 2022).

### Yeast Two-Hybrid (Y2H) and Bimolecular Fluorescence Complementation (BiFC) Assays

For Y2H, the full-length *FvM4K1*, phosphorylated site mutants, and different fragments of *FvM4K1* were cloned into pGBKT7, while *FvMOB1A* and *FvMOB1B* were cloned into pGADT7 (Lu et al., 2010b). Yeast strain Y2H Gold (Clontech) was used for transformation. Positive and negative control plasmids (pGBKT7-53/pGADT7-T and pGBKT7-Lam/pGADT7-T) were co-transformed into Y2H Gold as controls. Yeast cells were cultured in SD/−T/−L liquid media, and 10-fold serial dilutions were spotted onto selective SD/−T/−L and SD/−T/−L/−H/−A/AbA/X-α-Gal plates for reporter gene expression, and plates were scanned after four days. For BiFC, the same constructs used in Y2H were cloned into pGTQL1221YN/C (Lu et al., 2010b) and transformed into *Agrobacterium tumefaciens* strain GV3101 for infiltration into tobacco leaves. Confocal microscopy (Zeiss LSM 710) was used to observe fluorescence after infiltration. For DAPI staining, 5 µg/mL of DAPI was infiltrated into the leaves and incubated for two hours before observation under UV light.

### Subcellular Localization

Full-length FvM4K1 and FvM4K1-N1 (lacking the kinase domain) were cloned into pSuper1300 (Liang et al., 2020) to create C-terminal GFP fusion proteins. These constructs were transiently expressed in tobacco leaves and visualized by confocal microscopy.

### Protein Purification and *In Vitro* Kinase Assay

A truncated version of FvM4K1 lacking the first 236 amino acids (ΔN-FvM4K1) and its mutants (ΔN-FvM4K1^K269E^, ΔN-FvM4K1^T396A^, ΔN-FvM4K1^K269ET396A^) were cloned into the E. coli expression vector pET-28 (Novagen) and expressed in BL21 (DE3) cells. Protein expression was induced with 0.7 mM IPTG, and proteins were purified using His GraviTrap (Cytiva).

*In vitro* kinase assays were performed with 200 ng of purified ΔN-FvM4K1 or mutant proteins for auto-phosphorylation assays. To assess phosphorylation of FvMOB1 proteins or their mutants, 1000 ng of substrate protein was added to the reaction in kinase buffer (20 mM Tris-HCl, pH 7.5, 10 mM MgCl₂, 100 µM ATP, 10 mM EDTA, 10 mM NaF). The reaction was incubated at 30°C for 30 minutes, and proteins were resolved on a 10% SDS-PAGE gel. Phosphorylation was detected by Western blot using a phospho-threonine antibody (Cell Signaling), followed by ECL detection. Loading controls were assessed via EZBlue staining or Ponceau S staining.

### Data Analysis

Statistical analysis was performed using SPSS software. Error bars in the figures represent the standard deviation (SD), and the number of replicates is provided in the figure legends. A two-tailed t-test was used for statistical comparisons between two samples. For comparisons among multiple samples, one-way ANOVA followed by Duncan’s post hoc test was performed. Statistically significant differences (p < 0.05, as indicated in the figure legends) are denoted by different letters (e.g., a, b, c) or an asterisk (*).

### Accession Numbers

Sequence data for woodland strawberry in this study are available in the Rosaceae Genome database (RGD, https://www.rosaceae.org) and Arabidopsis in the Arabidopsis Biological Resource Centre (ABRC, http://www.arabidopsis.org/abrc/). The accession numbers are *FvM4K1* (FvH4_4g29800), *FvMOB1A* (FvH4_5g23830) and *FvMOB1B* (FvH4_1g08580) from RGD, *AtSIK1* (At1g69220), *AtMOB1A* (At5g45550) and *AtMOB1B* (At4g19045) from ABRC, *Ste20p* (Q03497), *Cdc15p* (P27636), *Hpo* (Q8T0S6), *hMst1* and *hMst2* (Q13043, Q13188) from Universal Protein Knowledgebase (Uniprot, https://www.uniprot.org/).

## Supplemental Data

**Table S1** Primers used in this study

