## Supplementary material for "FvM4K1, a Serine/Threonine Kinase and Ste20 Homolog Positively Regulates Fruit Size via Hippo Signalling Pathway in Woodland Strawberry (*Fragaria vesca*)": supplemantary figs

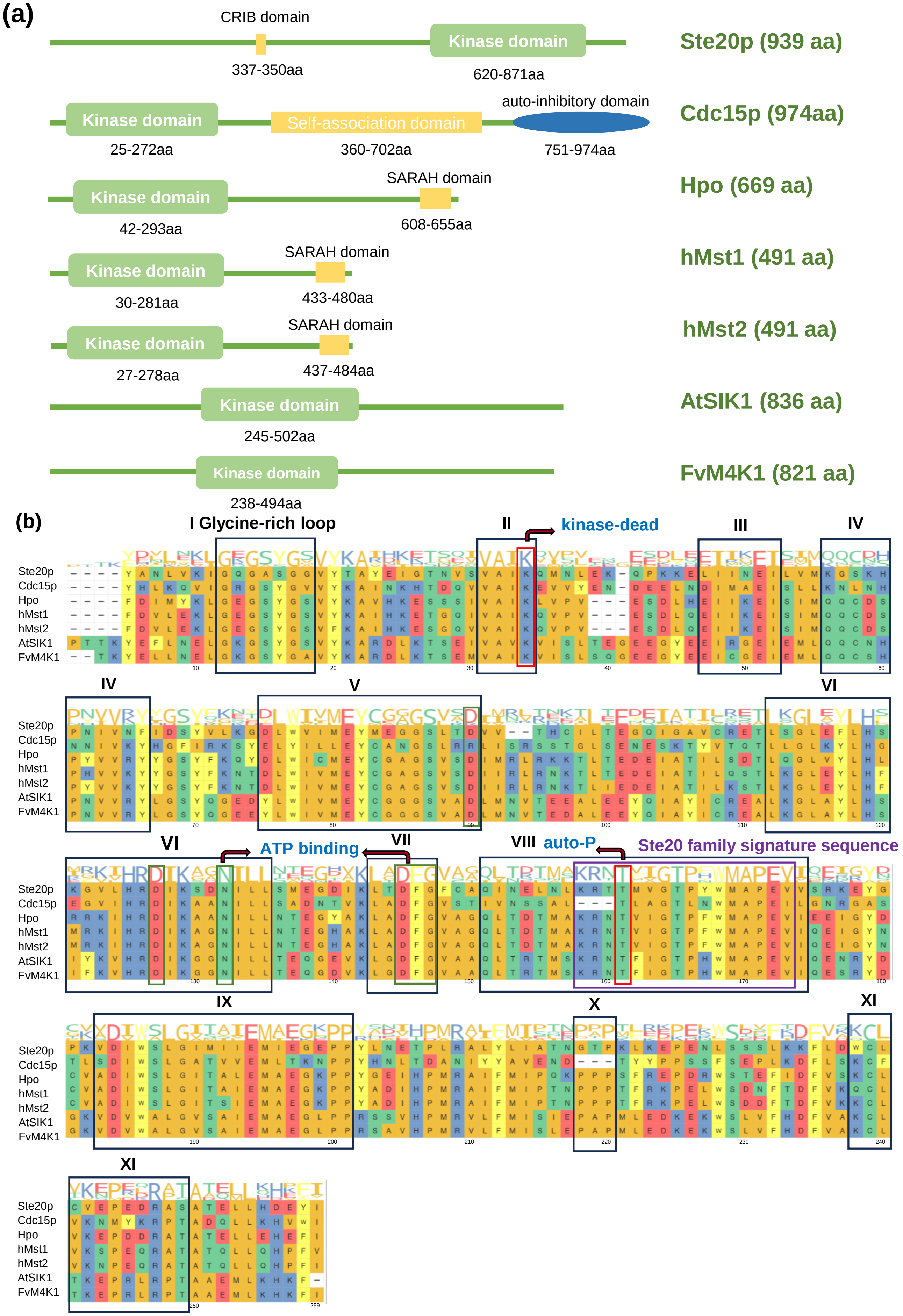


**Figure S1. Sequence comparison of the kinase domains between FvM4K1 of woodland strawberry and Ste20 proteins from Arabidopsis, yeast, fruit fly and humans.**

(a) Protein structure. The kinase domains of the Ste20 proteins are primarily located near the N-terminus, except for Ste20p which is close to the C-terminus. Only the kinase domain is found in FvM4K1 and AtSIK1 while others also have additional protein interaction domains.

(b) Protein sequence alignment. The consensus amino acids are shown at the top of the alignment. The larger capital letters of the amino acids are the more these residues are conserved among the Ste20 family proteins. Subdomains I to XI and conserved sites are boxed and labelled.

Accession numbers: Ste20p, YHL007c, Q03497, and Cdc15p, YAR019C, P27636 in yeast (*Saccharomyces cerevisiae*); Hpo, Hippo, Q8T0S6 in fruit fly (*Drosophila melanogaster*); hMst1/2, Mammalian Ste20-like protein kinase 1/2, Q13043, Q13188 in human; and Arabidopsis AtSIK1, At1g69220.

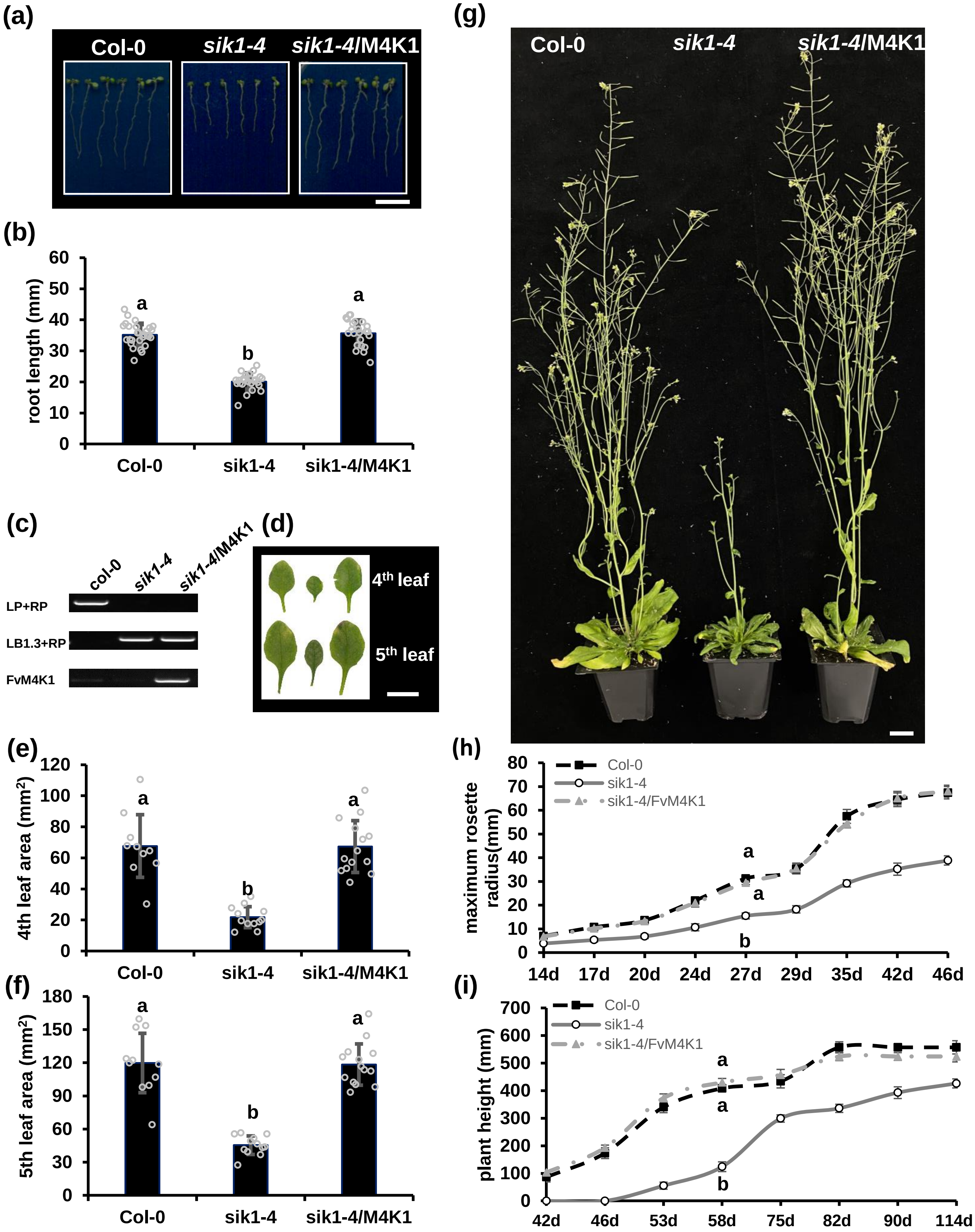


**Figure S2. FvM4K1 rescues the growth defects of Arabidopsis mutant *sik1-4*.**

(a) 10-day-old seedlings of Col-0 (WT), *sik1-4* (SALK_051369) and transgenic *sik1-4* expressing 35S:FvM4K1. Scale bar =10 mm.
(b) Root length of 10-day-old seedlings in (A). Error bars = means ± SD (n=20).
(c) Genotyping by PCR confirming the presence of *FvM4K1* in homozygous *sik1-4* but not in WT and *sik1-4* plants.
(d) Phenotypic analysis of the 4$^{th}$ and 5$^{th}$ leaves of 5-week-old plants of WT (left), sik1-4 (middle) and transgenic plant (right). Scale bar =10 mm.
(e) Area of 4$^{th}$ leaf. Error bars are means ± SD (n=10).
(f) Area of 5$^{th}$ leaf. Error bars are means ± SD (n=10).
(g) 8-week-old plants. Scale bar = 20 mm.
(h) Leaf maximum rosette radius. Error bars are means ± SD (n=20).
(i) Plant heights. Error bars are means ± SD (n=20).
Different letters above the columns in b, e, f, H&I indicate significant difference between samples at $p < 0.05$ calculated by one-way ANOVA with Duncan test using software SPSS.

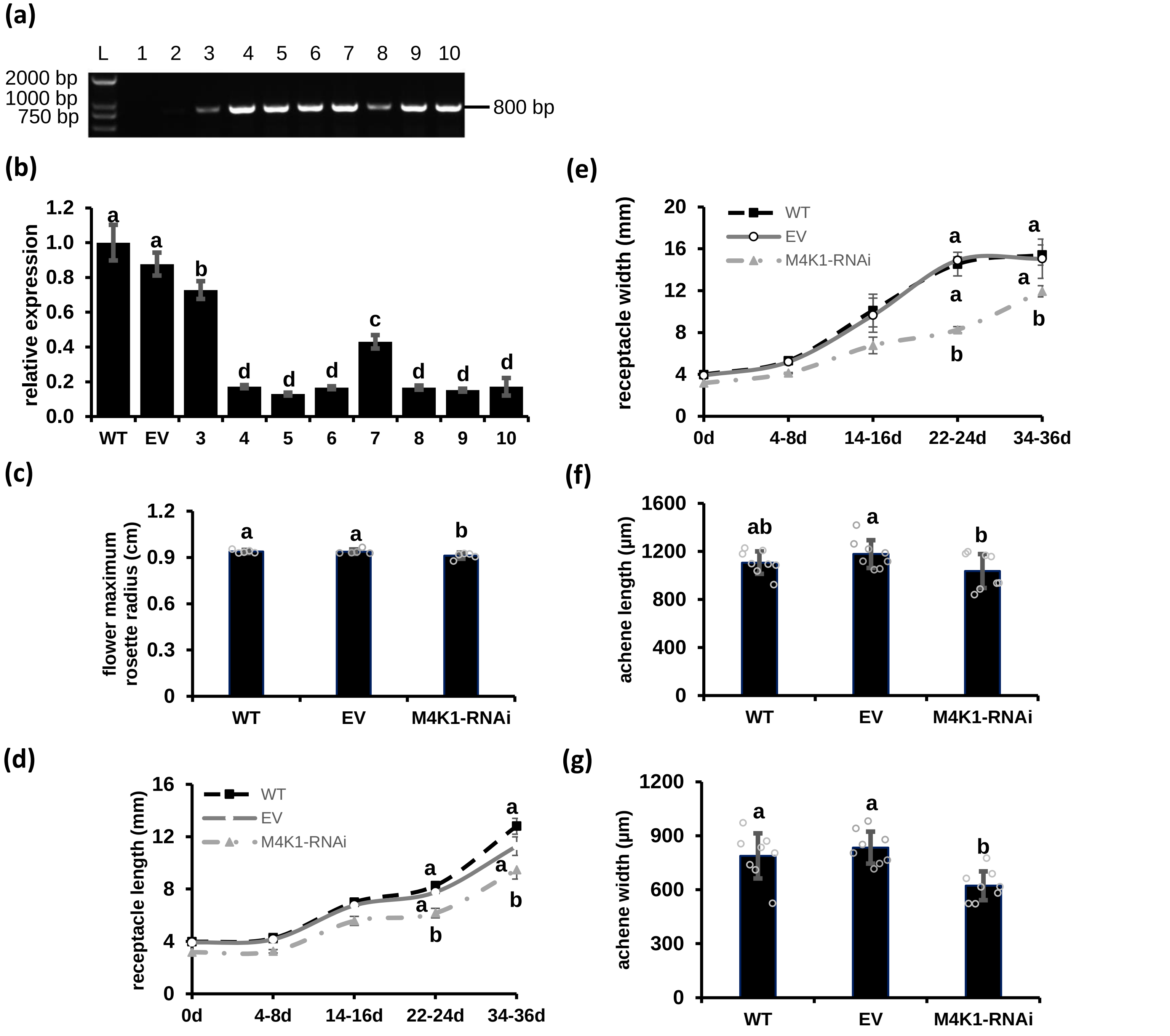


**Figure S3. Identification and observation of vegetative and reproductive organs of M4K1-RNAi transgenic plants.**

(a) Amplification of RNAi fragment (~800 bp) from genomic DNA isolated from 10 different lines of M4K1-RNAi strawberry plants, confirming the presence of the expression cassette in these plants. Lane 1-10, individual RNAi plants. The primer pair of 35S forward primer and CmR reverse primer (table S1) were used to amplify the DNA fragments.

(b) RT-qPCR to detect transcript level of *FvM4K1* in the WT, transgenic strawberry harbouring the empty vector (EV) and knock-down transgenic strawberry M4K1-RNAis. Lanes 3-10 were corresponding to lane 3-10 in (a). *FvActin* (FvH4_7g22410) was used as the reference gene. The relative expression of *FvM4K1* was calculated by the $2^{-\Delta\Delta Ct}$ method using *FvM4K1* expression level in WT as 1. At least three replicates were included in each run.

(c) Measurements of flower maximum rosette radius.

(d-e) Measurements of the length, width of the receptacles (fruits) at different developmental stages.

(f-g) Measurements of the length, width of the achenes in mature achenes (seeds).

Error bars are means ± SD (n=6). Different letter indicates statistically significant difference between RNAi, WT and EV plants at $p < 0.05$ calculated by one-way ANOVA with Duncan test using software SPSS.

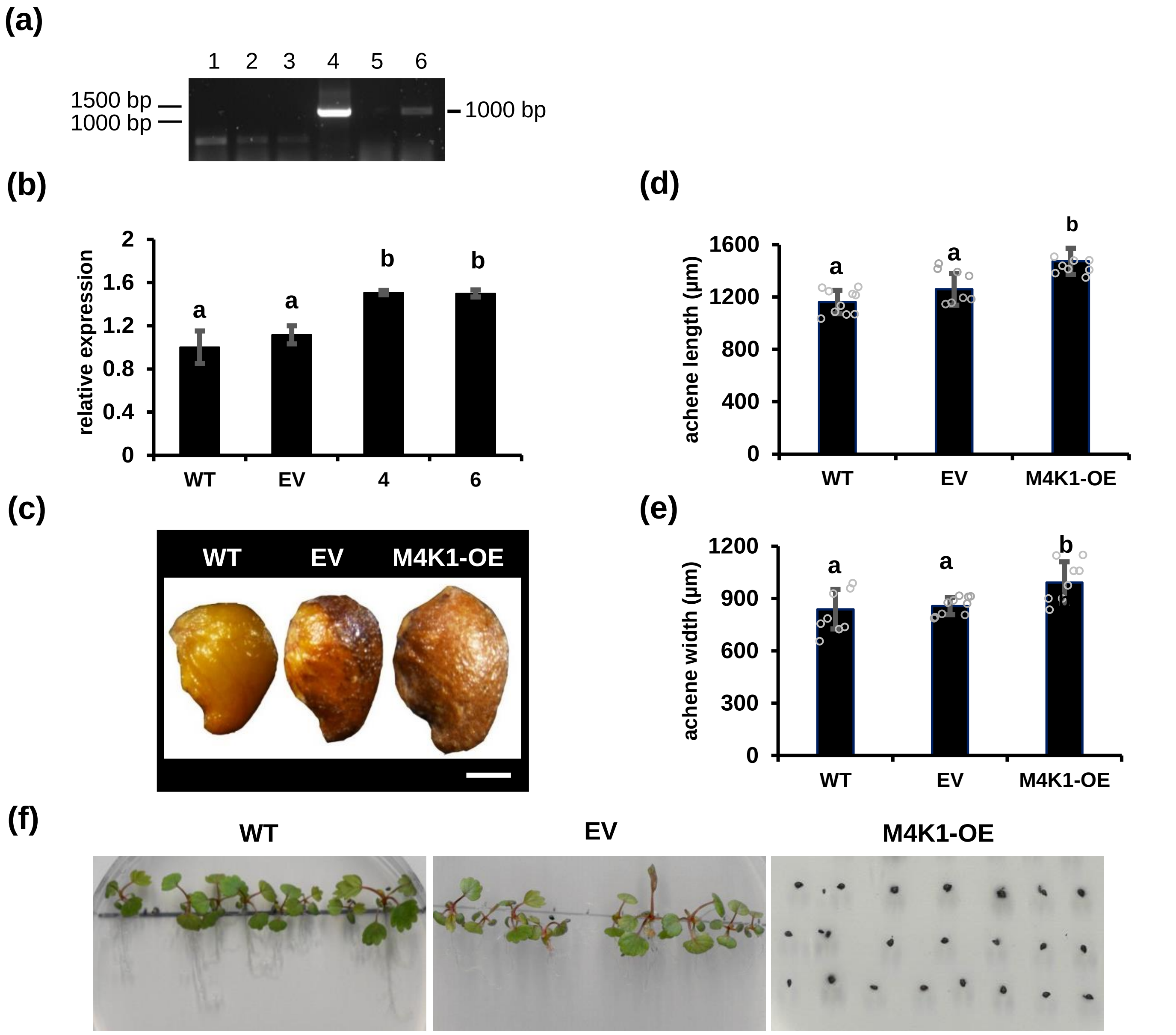


**Figure S4. Identification and observation of vegetative organs and achenes (seeds) of M4K1-OE transgenic plants.**

(a) Amplification of *M4K1* fragment (~1,000 bp) from genomic DNA isolated from 6 different lines of M4K1-OE strawberry plants. Lane 1-6, individual OE plants. The primer pair of 35S forward primer and gene specific reverse primer (table S1) were used to amplify the DNA fragments.
(b) RT-qPCR to detect transcript level of *FvM4K1* in the WT, transgenic strawberry harbouring the empty vector (EV) and overexpression transgenic strawberry M4K1-OEs. Lane 3, 4 were corresponding to lane 4, 6 in (a). *FvActin* (FvH4_7g22410) was used as the reference gene. The relative expression of *FvM4K1* was calculated by the $2^{-\Delta\Delta Ct}$ method using *FvM4K1* expression level in WT as 1. Three replicates were included in each run.
(c) Achenes (seeds) of WT, EV and M4K1-OE. Scale bar = 0.5 mm.
(d-e) Measurements of the length, width of the seeds (achenes) in mature achenes.
(f) After surface sterilization and vernalization, seeds (achenes) were placed on the ½ MS plate containing 1 % sucrose under long-day conditions for 35 days.

Error bars are means ± SD (n=) in (d-e). Different letter indicates statistically significant difference between RNAi, WT and EV plants at $p < 0.05$ calculated by one-way ANOVA with Duncan test using software SPSS.

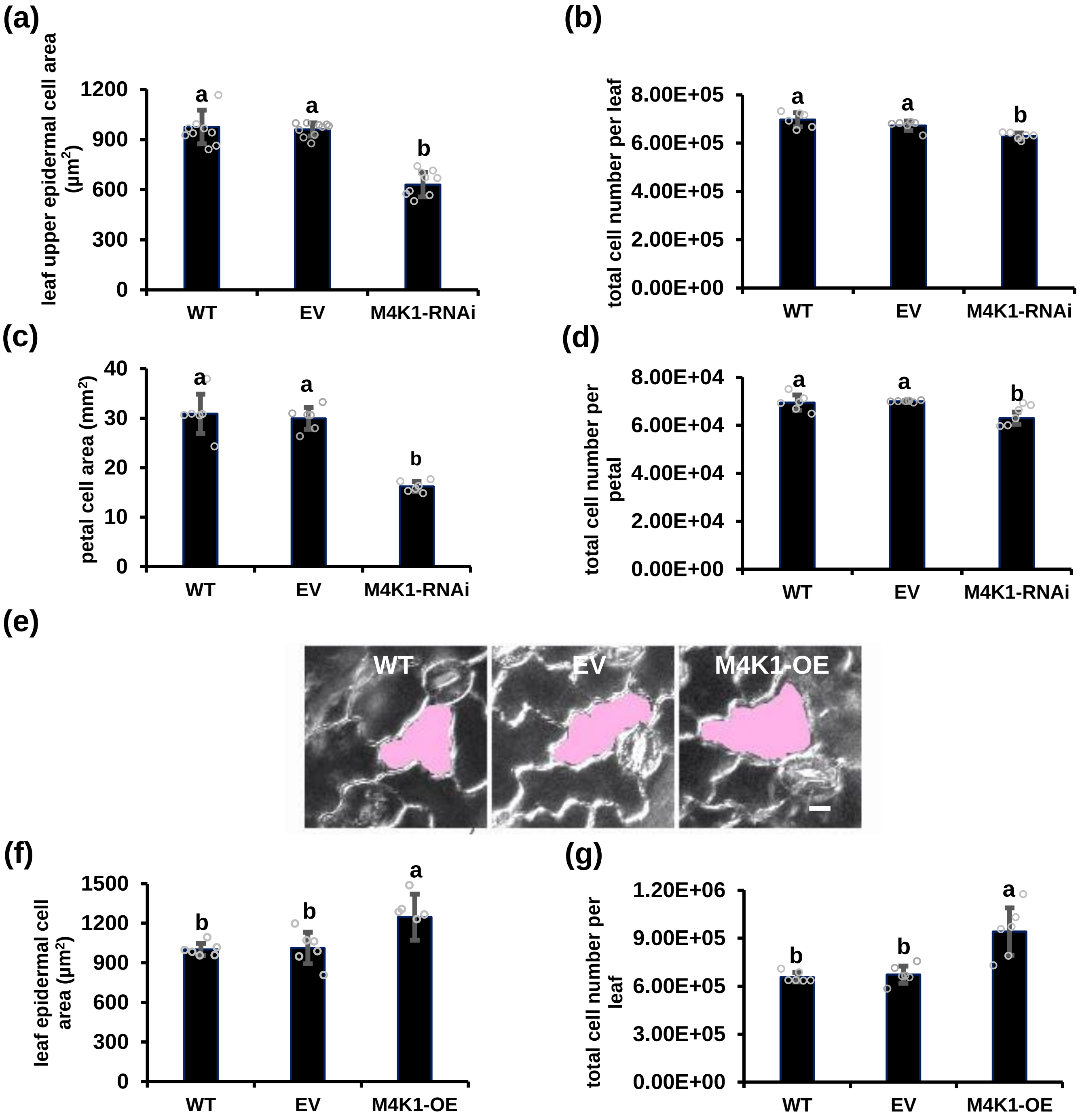


**Figure S5. The FvM4K1-RNAi knock-down plants have reduced cell size and number in the leaves and petals while M4K1-OE have increased in leaves.**

(a, b) Epidermic cell area of leaves and epidermic cell number per leaf from WT, EV and M4K1-RNAi plants.
(c, d) Epidermic cell area of petals, epidermic cell number per petal of WT, EV and M4K1_RNAi plants.
(e) The epidermal cells of leaves from WT, EV and M4K1_OE plants. A single cell from each genotype was shaded in pink to highlight their size difference. Bar = 10 μm.
(f, g) Epidermic cell area of leaves, epidermic cell number per leaf from WT, EV and M4K1_OE plants.

Error bars are means ± SD (n=6). Different letter indicates statistically significant difference between different plants at $p < 0.05$ calculated by one-way ANOVA with Duncan test using software SPSS.

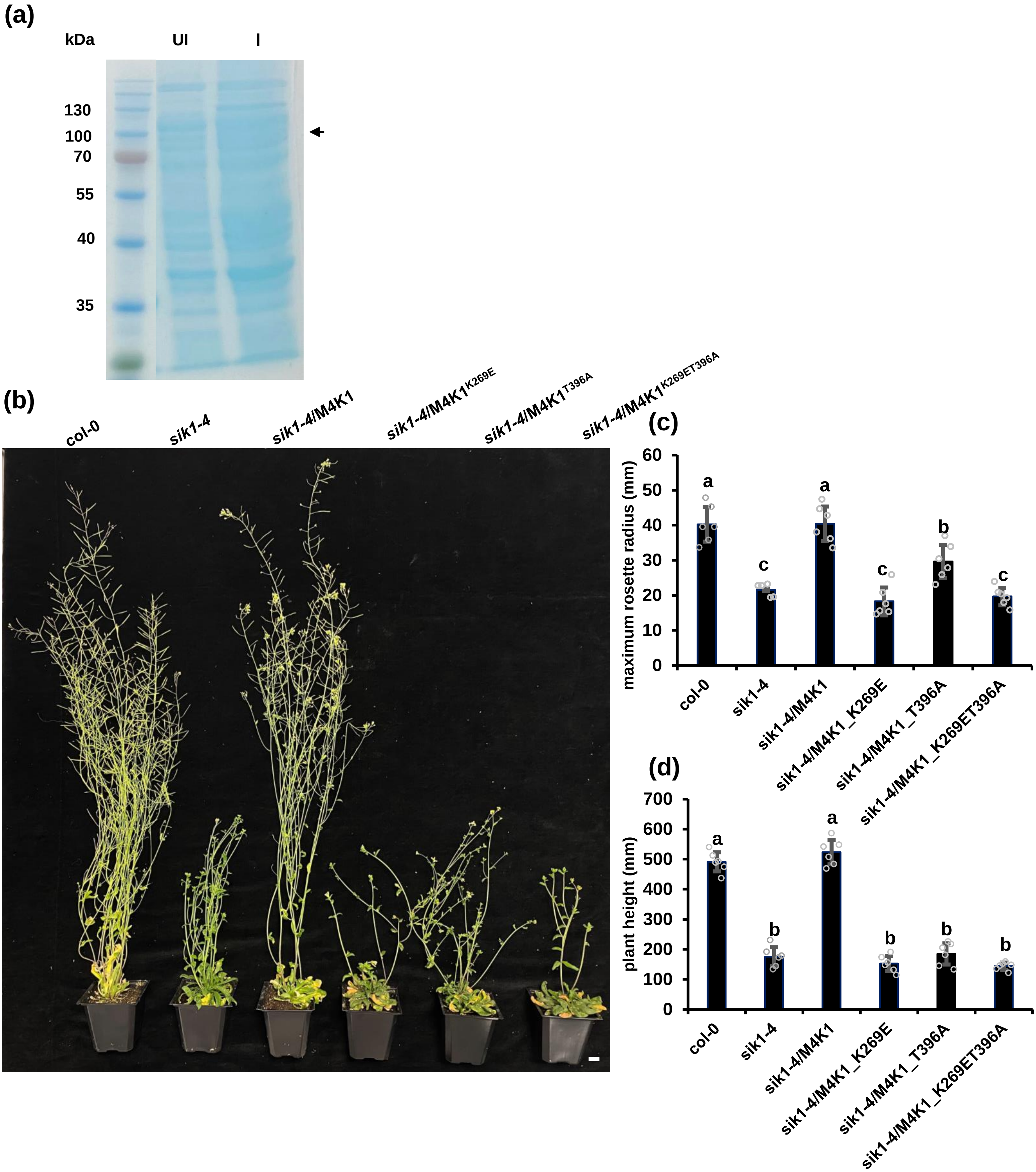


**Figure S6. FvM4K1 is auto-phosphorylated and its kinase activity is required for organ size control.**

(a) Full length of FvM4K1 cannot be expressed in BL21(DE3). UI, uninduced, I, induced with 0.7 mM IPTG at OD600 = 0.7 followed by incubation at 28°C for 4 hours. Black arrows indicate the predicted positions of FL-FvM4K1.

(b) 9-week-old Arabidopsis plants of Col-0, *sik1-4* and *sik1-4* expressing FvM4K1, FvM4K1$^{K269E}$, FvM4K1$^{T396A}$ and M4K1$^{K269ET396A}$. Scale bar =10 mm.

(c) Leaf maximum rosette radius and plant height. Error bars are means ± SD (n=6). Letters above the columns indicate statistically different between transgenic Arabidopsis expressed by FvM4K1 and its pointed mutants at $p < 0.05$ calculated by one-way ANOVA with Duncan test using SPSS.

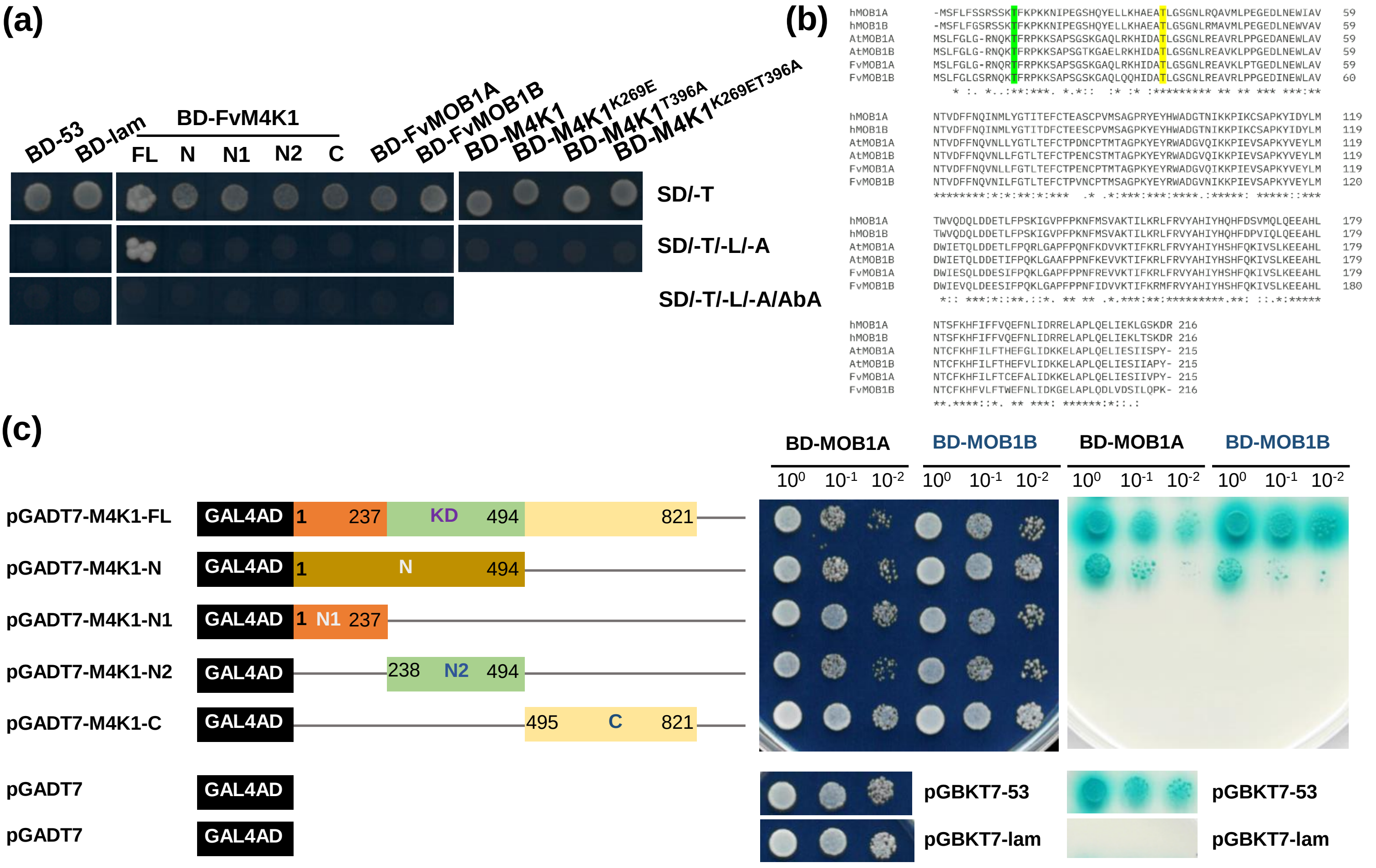


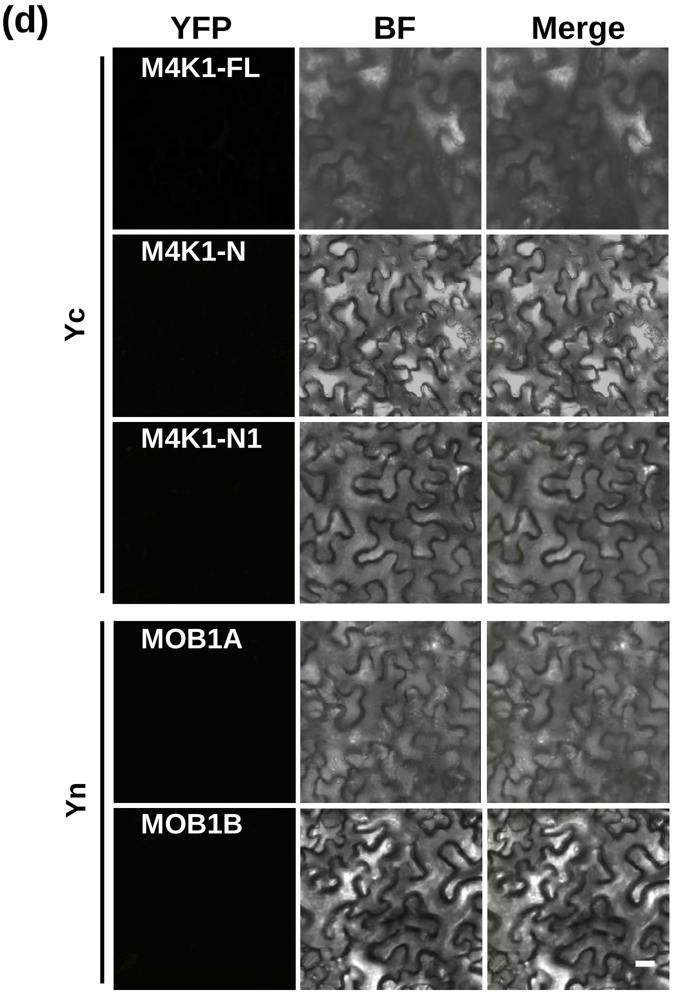


**Figure S7. FvM4K1 interacts with FvMOB1s in the Hippo pathway.**

(a) Self-activation test. BD-53 and BD-lam are positive and negative controls, respectively. SD/-T, SD/-T/-L/-A is nutrient selective media and SD/-T/-L/-A/AbA nutrient selective media containing 200 ng/mL of Aureobasidin A (AbA).
(b) Protein sequence alignment of hMOB1s, AtMOB1s and FvMOB1s. The conserved Thr12/13 and Thr35/36 were shaded green and yellow, respectively.
(c) FvM4K1 and its different fragments fused with AD interacted with FvMOB1A and FvMOB1B fused with BD in Y2H. '+', positive control pair pGBKT7-53 and pGADT7-T, '–', negative control pair pGBKT7-Lam with pGADT7-T. All colonies were grown to reach OD600=1, dilutions of 1/10, 1/100 were spotted on the plates. Colonies on the left panels grew on the media SD/-T/-L while on the right on SD/-T/-L/-H/-A/AbA/X-α-Gal containing AbA (200 ng/mL) and X-α-Gal (40 µg/mL).
(d) BiFC controls. FvM4K1 and its different fragments fused with neYFP and empty vector ceYFP on the left panel, FvMOB1A&B fused with ceYFP and empty vector neYFP.

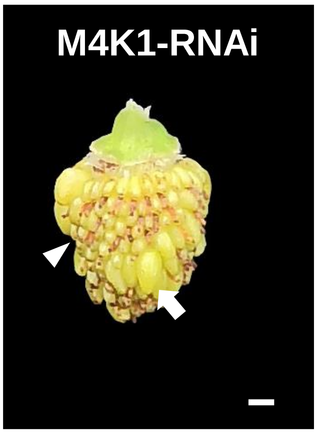


**Figure S8. Defect of receptacles (fruits) of M4K1-RNAi.**

Observation of achenes (seeds) of M4K1-RNAi at 15 DAF. White arrow indicates developing seed, white triangle indicates aborted seeds. Scale bar = 5 mm.

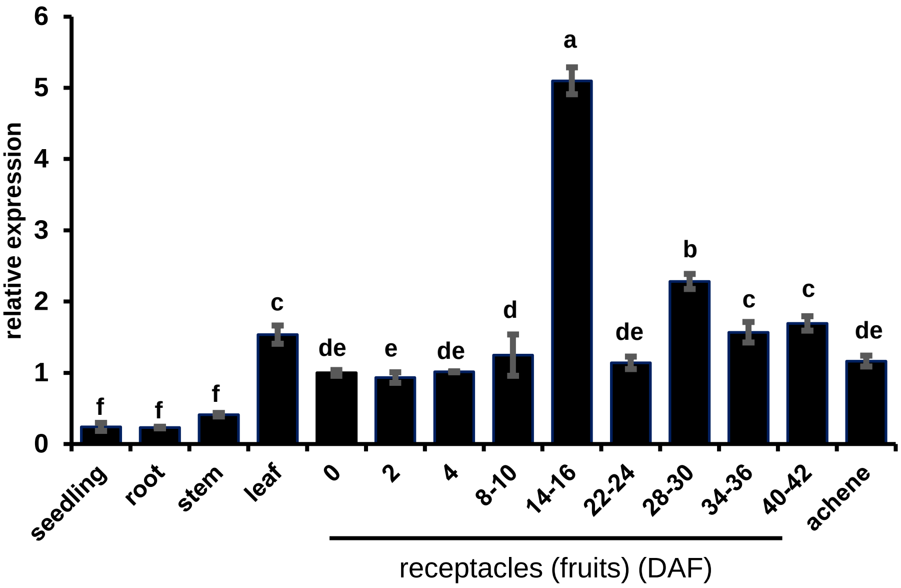


**Figure S9. Expression profile monitored by RT-qPCR of *FvM4K1* in different tissues and receptacles at different developmental stages.**

The transcripts of *FvM4K1* in 14-day-old seedlings, roots, stems, leaves, achenes of mature plants, as well as receptacles at 0, 2, 4, 8-10, 14-16, 22-24, 28-30, 34-36, 40-42 day-after-flowering (DAF). The relative transcript level of *FvM4K1* is calculated by the $2^{-\Delta\Delta Ct}$ method using *FvActin* (FvH4_7g22410) as the reference gene and the *FvM4K1* expression level of flowers (0 DAF) as 1. Error bars = means ± SD (n=3). The different letters indicate significant difference between samples at $p < 0.05$ calculated by one-way ANOVA with Duncan test using SPSS.
